# Multi-modal profiling of biostabilized human skin modules reveals a coordinated ecosystem response to injected mRNA-1273 COVID-19 vaccine

**DOI:** 10.1101/2023.09.22.558940

**Authors:** Manon Scholaert, Mathias Peries, Emilie Braun, Jeremy Martin, Nadine Serhan, Alexia Loste, Audrey Bruner, Lilian Basso, Benoît Chaput, Eric Merle, Pascal Descargues, Emeline Pagès, Nicolas Gaudenzio

## Abstract

The field of vaccination is witnessing a remarkable surge in the development of innovative strategies. There is a need to develop technological platforms capable of generating human data prior to progressing to clinical trials. Here we introduce VaxSkin, a flexible solution designed for the comprehensive monitoring of the natural human skin ecosystem’s response to vaccines over time. Based on bioengineering to repurpose surgical resections, it allows a comprehensive analysis of the response to vaccines at both organ and single-cell levels. Upon injection of the mRNA-1273 COVID-19 vaccine, we characterized precise sequential molecular events triggered upon detection of the exogenous substance. We also found that the vaccine consistently targets DC/macrophages and mast cells, regardless of the administration route, while promoting specific cell-cell communications in surrounding immune cell subsets. Given its direct translational relevance, VaxSkin provides a multiscale vision of skin vaccination that could pave the way toward the development of new vaccination development strategies.

## Introduction

The human immune system is composed of a plethora of specialized cell types distributed across organs and modulated throughout a person’s life upon exposure to environmental triggers. This unique, adaptable feature of tissue-resident immune cells contributes to the establishment of immunological memory for each individual and to drive a coordinated and immediate defense against pathogenic stimuli^1^. The skin is one of the largest organs of the body and is composed of a wide variety of myeloid cells (e.g., dendritic cells (DCs), macrophages, Langerhans cells (LCs), mast cells, etc.) and lymphoid cells (e.g., CD4 T cells, CD8 T cells, etc.) that interact with an organized network of structural elements (e.g., keratinocytes, stromal cells, blood vessels, etc.) to form a tightly-regulated ecosystem^2^. Such unique immune features of the skin make it a promising anatomical site for studying the human immune response *ex vivo*, while preserving the natural complexity at the organ level.

Recent advances in mRNA vaccine technology have led to the development of highly effective vaccines, which utilize lipid nanoparticles (LNP) to deliver the mRNA into cells. These vaccines have demonstrated remarkable efficacy in clinical trials^3,4^ and have been authorized for emergency use worldwide during the COVID-19 pandemic^5^. With the increasing number of academic laboratories, biotechnology, and pharmaceutical companies developing new mRNA-based vaccines and exploring alternative routes of administration through the skin^6,7^, there is a critical need for scalable technological platforms to evaluate the immunogenicity and safety of vaccines before advancing to clinical trials. Notably, how human tissue-resident structural and immune cells interact with, and are modulated by, mRNA-loaded LNPs at the site of injection remains a promising area of investigation^8^.

Current *in vitro* technologies have made significant strides in replicating selected aspects of the human skin as a tissue; however, they remain unable to capture the complexity of the native organ and the genetic diversity found in the human population^9–11^. For example, microphysiological systems such as organ-on-a-chip technologies^12^, skin substitutes^13^ and bioprinting approaches^14^ can mimic important cellular and drug permeability features, yet they lack the multifaceted 3D architecture, the cellular interactome and, most importantly, the inherent complexity of the human immune system. The accessibility to large quantities of donated human skin leftover from surgical resections has prompted scientists to use cultured skin punch biopsies to better understand the response to vaccines^15,16^ . The existing limitations of current *in vitro* culture protocols make it challenging to preserve the viability of skin explants for extended periods of time, in particular the subcutaneous adipose tissue, which is an essential component for subcutaneous administration^17,18^. Additionally, classical floating culture conditions can lead to the escape of immune cells into the culture medium, directly affecting the accuracy of the results under investigation^19,20^.

Here we describe the VaxSkin platform, a general framework primarily designed for the longitudinal profiling of the human skin ecosystem in response to vaccines at the site of injection. We first engineered a chemically-defined jellified matrix that maintains the viability and immunocompetence of the skin for a period of 7+ days and coupled to a topical silicone ring that preserves its tension *ex vivo*^21^. Each individual human skin “module” (i.e., association of matrix-skin biopsy-silicone ring) can be injected subcutaneously or intradermally with a conventional clinical syringe, as the system maintains the integrity of the native adipose tissue attached to the dermis. The method is scalable enough to conduct multiple parallel experiments, including transcriptomics, 3D imaging, and secretomics in one donor and then to extend to selectable cohorts based on age, gender, and ethnicity. We designed an adaptable analytical pipeline composed of integrated multiparametric analyses to extract complementary immune datasets from each individual module, including an *in silico* approach to selectively track and quantify mRNA vaccines incorporation at the single cell level.

## Results

### Biostabilized human skin modules are structurally stable and injectable subcutaneously and intradermally

The preservation of skin viability is one of the major challenges in developing *ex vivo* models that accurately mimic the human organ outside of the body. To circumvent this limitation, we developed a series of standardized engineering steps to quickly biostabilize human skin biopsies, while preserving an air-liquid interface and the appropriate biomechanical properties of the tissue. Upon surgical excision from donors, large skin biopsies (i.e., an average size of 365 cm²) were punched into individual skin explants of 23 mm diameter using a precision mechanical press in a controlled laboratory environment. Up to 50 individual explants can be generated from one individual donor. Subsequently, we encased each explant into a solid gel matrix and a silicone ring was placed on the top to emulate natural tension and maintain the biomechanical property found in the skin. These individual human skin “modules” were then carefully positioned in a transwell system and bathed in a chemically-defined culture medium devoid of animal product, which was replaced on a daily basis (**Fig. 1a**).

To evaluate potential donor-intrinsic changes in global tissue architecture and cellular gene expression profiles over the culture period, we performed a combination of histochemical and single cell RNA sequencing (scRNAseq) longitudinal analyses, within the same individual donor (**Fig. S1, upper panel**). We first used hematoxylin and eosin (H&E) staining, a commonly used method in histology to visualize most tissue structures. No apparent changes were observed in the integrity or overall cellular structure of the skin over 7 days. The epidermal and dermal layers remained visually unaltered, with preserved keratinocytes and collagen fibers, respectively. We could detect some discrete morphological changes in the epidermis-dermis junction and a tendency for dermal fibroblast to decrease in number when the skin modules were analyzed at day 10. Interestingly, the subcutaneous adipocyte layers remained morphologically unaltered over the 10-day culture period (**Fig. 1b**). Using fluorescence microscopy, we analyzed the expression of active caspase-3, as a readout of tissue apoptosis, and the distribution of filaggrin and keratin 14 to assess changes in the distribution of key epidermal structural proteins^22^. The active caspase-3 was detected in few cells at day 0 and its expression did not increase over the culture period (**Fig. S2a**). The filaggrin and keratin 14 were expressed in the stratum corneum and the basal layers, respectively, during the entire culture period (**Fig. S2b, c**). We next used two-photon imaging to assess the stability of the following additional skin components at the protein level in a large field of view and in 3D: claudin 1 (a key component of the epidermal barrier), CD45 (an immune cell marker), CD31 (a blood vessel marker), and β3-tubulin (that identifies peripheral nerve endings). In line with our previous observations (**Fig. 1b, S2**), all analyzed fluorescent markers were detected in the skin modules following 7 days of culture, without noticeable changes in either intensity or anatomical distribution (**Fig. 1c, Videos 1 and 2**).

To explore the suitability of the human skin modules as a model for studying injectable drugs, we conducted subcutaneous (s.c.) and intradermal (i.d.) infusion experiments using a clinical grade syringe. We introduced a colored aqueous solution directly into the adipose layer (s.c.) or the dermis (i.d.) of the skin within the modules. This allowed us to examine the distribution and diffusion patterns of the injected solutions, providing insights into the potential leakage of injectable drugs out of the tissue. We found that 60 minutes after a single s.c. injection, the colored solution remained within the subcutaneous compartment, and started to slowly diffuse in the upper layers of the skin, including the dermis and epidermis (**Fig. 1d, right panel**). Conversely, after a single i.d. injection, the colored solution remained within the dermal compartment, and started to diffuse slightly into the epidermis and the subcutaneous space (**Fig. 1d, left panel**). These data were also confirmed using 3D X-ray tomography analysis after the injection of a radiocontrast agent (**Fig. 1e** and **Videos 3 and 4**). Importantly, there was no detectable leakage of the solution outside of the skin explant, indicating a well-contained distribution of the injected substance within the module environment.

**Fig. 1.**
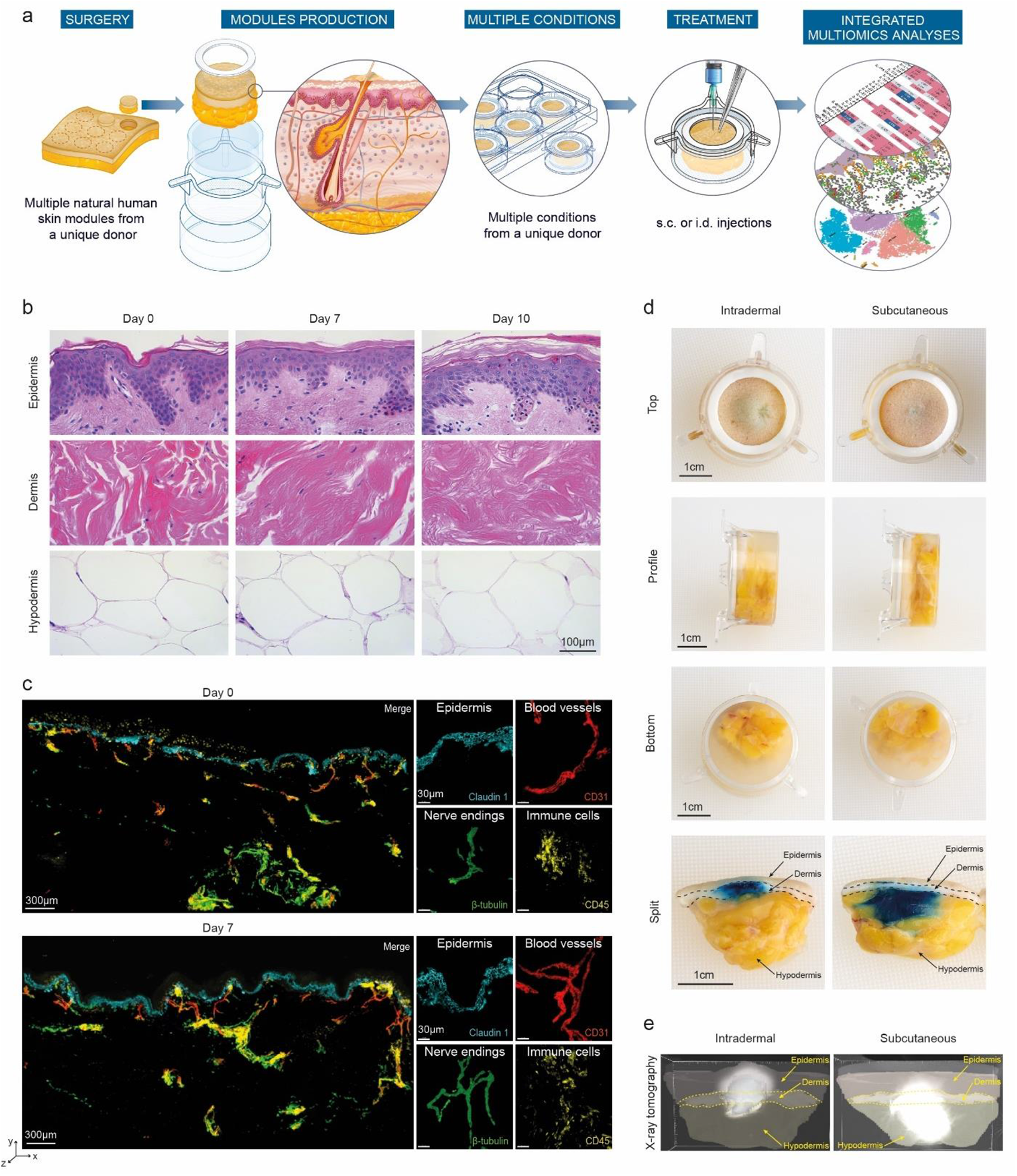
Biostabilization and injectability of natural human skin modules. **a**, Schematic for e*x vivo* natural human skin modules bioengineering process and analytical pipeline, from production using skin leftovers from surgical procedures to the extraction of human biological datasets using scaled multiomics approaches. **b**, Comparison of H&E staining of natural human skin modules after 0, 7 and 10 days of culture. **c**, Representative 3D two-photon image of immune cells (yellow), epidermis (cyan), blood vessels (red), and nerve fibers (green) of natural human skin modules after 0 and 7 days of culture. **d**, Intradermal (left panel) or subcutaneous (right panel) injection of blue dye in natural human skin modules. Top, profile, bottom and split views of the model are presented. **e,** 3D representation acquired by X-ray tomography, showing the bolus in the dermis (left panel) or in the hypodermis (right panel), respectively. Bars = 100 μm (B), 300 μm (C) and 1 cm (D). *s.c: subcutaneous; i.d: intradermal*.

### Longitudinal assessment of the transcriptomic program of human skin modules at the single cell level

We next investigated potential subtle transcriptomic changes in phenotype and/or activation status of both structural and immune cells overtime. We produced human skin modules of 23 mm diameter from the same individual and performed 10x Genomics scRNAseq analysis of the dissociated skin at day 0 (i.e., within 24h after surgery), 3, 5 and 10 (**Fig. S1, upper panel**) to generate a longitudinal transcriptomic profile for each individual cell (all quality controls are described in the Methods section). Using the t-distributed stochastic neighbor embedding (t-SNE) approach, we analyzed the expression patterns of 36,601 genes in 26,553 individual cells from the aggregated dataset of all time points and could unambiguously identify the presence of 9 global clusters (**Fig. 2a**). All populations were annotated based on their combined expression of cardinal marker genes, such as *KRT14* (undifferentiated keratinocytes), *KRT10* (differentiated keratinocytes), *KRT17* (proliferating keratinocytes), *SERPINB4* (inflammatory keratinocytes), *LYVE1* (lymphatic endothelial cells, LECs), *ACKR1* (vascular endothelial cells, VECs), *COL6A1* (fibroblasts), *PTPRC* and *CD3E* (lymphoid cells), *PTPRC* and *CD1A* (myeloid cells), and *MLANA* (melanocytes) (**Fig. 2b**). We next analyzed the distribution of all cells on the aggregated t-SNE graph based on their time point of analysis. We observed an overall homogeneous distribution of the cells on the aggregated t-SNE based on their original phenotype, but not based on the time point of analysis (**Fig. 2c**). In order to objectively evaluate any transcriptomic variations at the tissue level, we applied a pseudo-bulk transformation in the global single-cell dataset of each time point^23^. By calculating a Pearson correlation heatmap, we assessed the correlation between the transcriptomic profiles of day 0, 3, 5, and 10. We observed a consistently high correlation exceeding 0.995 across all time points, indicating that the overall transcriptomic program of the skin remained stable over time (**Fig. 2d**). In line with these findings, we found that all identified clusters harbored similar proportions of cells in either G1, S, or G2M phases, suggesting a conserved cell cycling capability in each population (**Fig. 2e**).

The human skin undergoes continuous turnover of keratinocytes, ensuring the constant renewal of the epithelial barrier. These cells undergo a process of differentiation, transitioning from basal cells in the innermost layer of the epidermis to terminally differentiated cells in the outermost layer. This process involves sequential changes in gene expression, leading to the production of specific proteins (including keratins) and the formation of a protective barrier against external factors^24^. We observed a thickening of the cornified layer over the culture period in the absence of mechanical exfoliation (**Fig. 1b**), which suggests a continuous turnover of keratinocytes in the human skin module environment. We further tested this hypothesis by using *Monocle3* pseudotime analysis^25^ on keratinocyte single-cell datasets to reconstruct the temporal order of cellular states and identify the progression of differentiation over time.

We found that the generated pseudotime trajectory recapitulated the normal differentiation of keratinocytes through the following consecutive states: undifferentiated, proliferating, inflammatory, and differentiated (**Fig. S3a-b**). In line with these findings, we could identify the progressive loss or acquisition of key genes characteristic of keratinocyte maturity states along the pseudotime axis, such as *KRT14*, *ITGA6* and *ITGB1* (i.e., markers of immaturity), and *DSG1*, *KRT1* and *KRT10* (i.e., markers of maturity) (**Fig. S3c**).

Subsequently, we once again utilized a pseudo-bulk transformation method to assess the correlation between the transcriptomic profiles of day 0, 3, 5, and 10, but at the level of individual populations to potentially detect subtle changes over time. We observed a high correlation of more than 0.985 across all time points for melanocyte, myeloid, lymphoid, and keratinocyte populations. However, slightly lower correlations of 0.9710 and 0.9210 were found in endothelial cells and fibroblasts, respectively, after 5 days of culture of the skin modules (**Fig. 2f**). Finally, we applied the algorithm CellPhoneDB, a statistical framework coupled with a ligand-receptor interaction repository which allowed us the exploration of cell-cell communication within complex cellular ecosystems^26^. Through a comprehensive analysis of our datasets at each time point, we discovered that myeloid cells exhibited a significant capacity of interaction with a wide range of structural and immune cell types within the tissue, suggesting their potential pivotal role within the skin ecosystem (**Fig. 2g**). Consistently with our earlier correlation findings (**Fig. 2f**), we observed that endothelial cells displayed a propensity for intercellular communication with various cell types within the tissue, although this communication significantly decreased by day 10.

Taken together, these data demonstrate that the intercellular ecosystem naturally present in the *ex vivo* human skin modules remained relatively stable over 10 days of culture, at least at the microscopic and transcriptomic levels, although some variability seemed to emerge in the fibroblast and endothelial cell compartments. The prolonged stability observed in the immune compartment, particularly the myeloid cells, strongly supports the potential of natural human skin modules to be employed for comprehensive profiling of innate molecular mechanisms activated in response to vaccines at the site of injection.

**Fig. 2.**
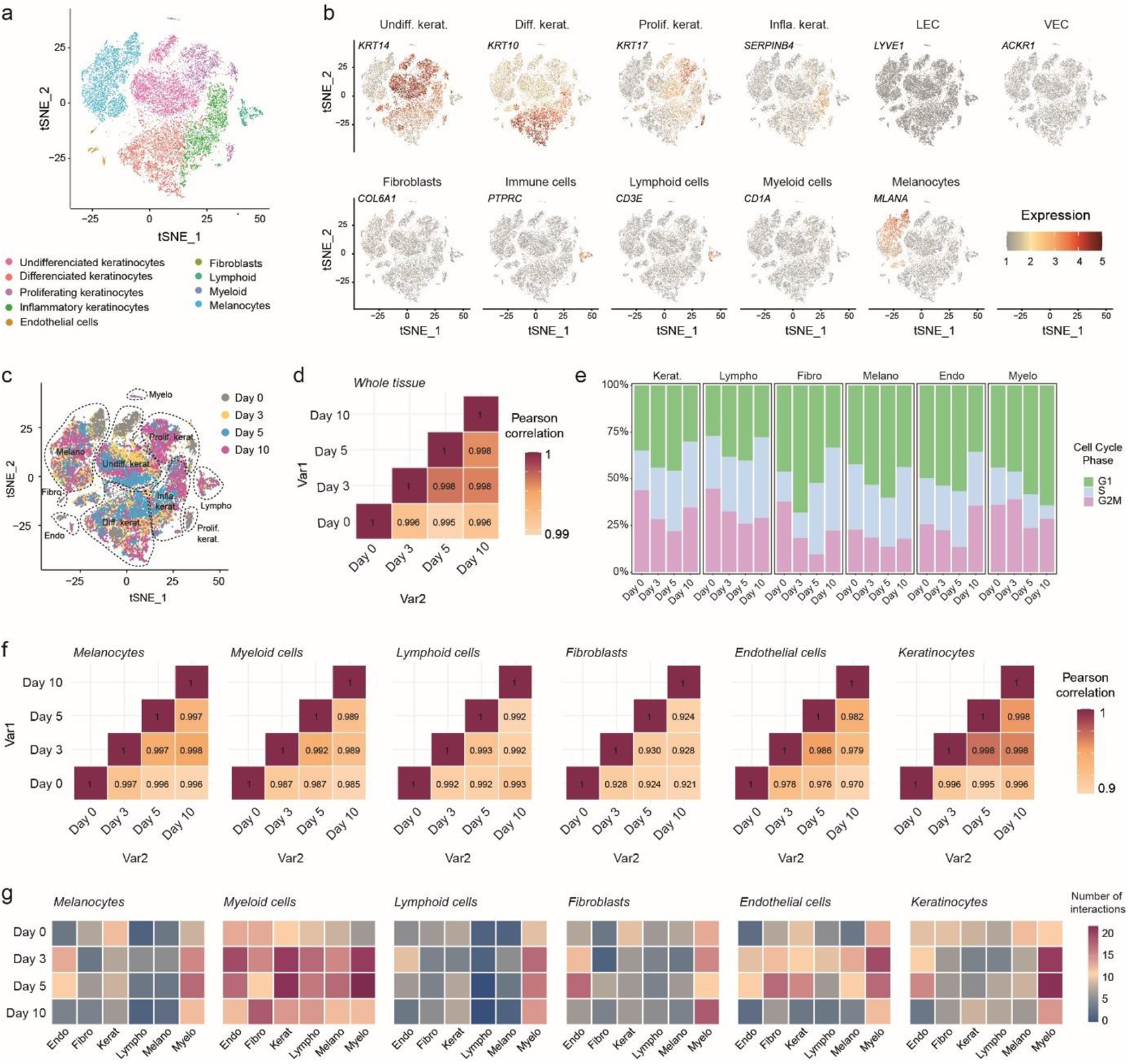
Transcriptomic stability of structural and immune cells over time in natural human skin modules. **a**, t-SNE plot of the scRNAseq aggregate performed after 0, 3, 5, and 10 days of culture and cell subsets attribution. **b**, Expression of cell-specific markers on the scRNAseq aggregate. **c**, t-SNE plot of the scRNAseq aggregate colored by timepoint. **d**, Pseudo-bulk correlation heatmap of the scRNAseq aggregate at different time points. The correlation coefficients between each pair of timepoints are indicated in the correlation map. **e**, Bar plot representing the repartition of the cell cycle phases for each cell type and day of culture. **f**, Pseudo-bulk correlation heatmap of each cell type and day of culture. The correlation coefficients between each pair of timepoints are indicated in the correlation map. **g**, Heatmaps of the number of interactions between all cell types at different days of culture. *Undiff. kerat.: Undifferentiated keratinocytes; Diff. kerat.: Differentiated keratinocytes; Prolif. kerat.: Proliferating keratinocytes; Infla. kerat.: Inflammatory keratinocytes; LEC: Lymphatic Endothelial Cells; VEC: Vascular Endothelial Cells; Lympho: Lymphoid cells; Myelo: Myeloid cells; Melano: Melanocytes; Fibro: Fibroblasts*.

### Quantitative and qualitative validation of reactogenicity in response to mRNA vaccination in human skin modules

Vaccines are often developed using a combination of congenic mouse experiments and *in vitro* culture of human peripheral blood cells. While such experimental settings are still very useful, they often fail to recapitulate the complexity of the human immuno-structural ecosystem found in the tissue at the site of injection^2^. With this in mind, we decided to benchmark the effective performance of the *ex vivo* human skin modules as a proxy to understand subtle cellular and molecular changes in response to subcutaneous infusion of the commercially-available Moderna mRNA-1273 COVID-19 vaccine^27^ since this vaccine is typically administered into the muscle through the skin and subcutaneous tissue. This allowed us to investigate and gain insights into the potential local effects of the vaccine on immune cells and structural components, at both the tissue and single-cell levels. To achieve this, we implemented VaxSkin, a flexible analytical pipeline composed of integrated multi-modal analyses specifically adapted to extract qualitative and quantitative information from the biostabilized skin explants with a diameter of 23 mm (**Fig. S1, lower panel**).

We injected the vaccine in natural human skin modules generated from 6 different donors and measured 36 cytokines and chemokines released from the injected tissue in the culture medium after 8h and 24h using a electrochemiluminescence assay from Meso Scale Diagnostics^28^. When compared to control modules from the same donors injected with clinical-grade water, following vaccine administration, we observed a general increase the levels of interleukin (IL)−7, IL−12/IL−23p40, IL−15, IL−16, IL−1α, IL−8(HA), IL−5, IL−17A, vascular endothelial growth factor (VEGF), macrophage inflammatory protein (MIP)−1α, MIP−1β, granulocyte-monocyte colony-stimulating factor (GM−CSF), interferon gamma-induced protein (IP)−10 (also known as CXCL10), human macrophage/monocyte chemotactic protein (MCP)−4, and Eotaxin. The following cytokines were not detected under our experimental conditions: TNF-β, IL-31, IL-27, IL-23, IL-22, IL-21, Eotaxin 3, and IL-17A Gen B. Conversely, 4 cytokines and chemokines known to be secreted by activated T cells, DCs, LCs, and macrophages were found to be significantly increased. This conserved secretomic signature triggered by the injection of the vaccine across the 6 donors tested includes IFN-γ, the thymus and activation-regulated chemokine (TARC, also known as CCL17), MCP−1 (also known as CCL2)^29^, and the macrophage-derived chemokine (MDC, also known as CCL22)^30^ (**Fig. 3a**). We then used this secretomic signature to infer which biological pathways could be regulated at the tissue level using a combination of protein-based publicly-available databases such as KEGG, GO-bp, and Reactome. We found a common signature of pathways associated with global chemotaxis, nitric-oxide biosynthesis, immune activation (e.g., Dectin-1-associated NF-κB activation or IL-17 signaling), signal transduction (e.g., C−type lectin receptor signaling pathway), nitric oxide synthase biosynthesis (i.e., innate defense mechanism that notably inhibits viral replication), and viral protein interaction with cytokines (**Fig. 3b** **and Table S1**).

Taken together, these findings strongly suggest that immune and/or structural cells, which naturally reside in natural human skin, maintain the ability to release immunomodulatory substances in response to vaccine administration in the modules. They also show that the injected vaccine induces the secretion of important chemokines known to play a role in APCs recruitment and vaccination effectiveness.

**Fig. 3.**
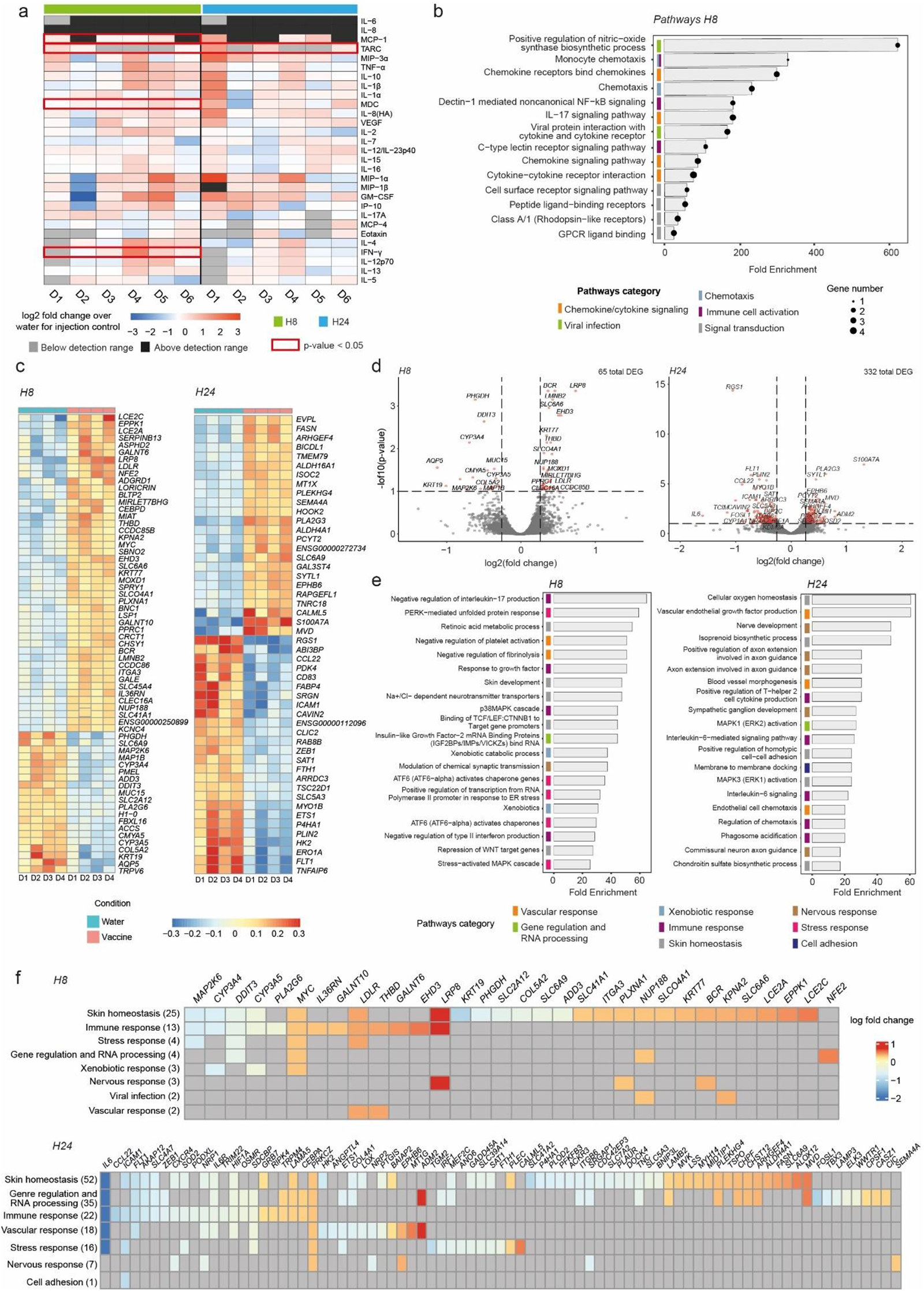
Activation of immune pathways following COVID-19 vaccine s.c. injection through cytokines secretion in natural human skin modules. **a**, Heatmap of the log2 fold change of the cytokines concentrations in clinical-grade water vs vaccine injected modules after 8h and 24h on 6 different donors. **b**, Barplot of the biological pathways associated with the differentially regulated cytokines 8h after injection, sorted by fold enrichment. **c**, Differentially-expressed genes (DEGs) after s.c. injection of clinical-grade water or COVID-19 vaccine at 8h (left panel) and 24h (right panel) in 4 different donors. At 8h, all the DEGs were presented and at 24h, the top 50 genes. **d**, Corresponding Volcano plots. **e**, Main biological pathways identified based on DEGs. **f**, Expression of selected genes associated with the biological pathways described in **e**.

We next performed bulk RNAseq analyses to characterize sequential changes in gene expression profiles at the tissue level, at 8h and 24h after s.c. administration of the COVID-19 vaccine or control clinical-grade water. As potential inter-individual variations and gender-based differences can introduce experimental biases, we applied a well-established *Combat-Seq* batch effect adjustment tool from R library *sva* version 3.48^31,32^. We next analyzed our datasets and detected 65 and 332 differentially expressed genes (DEGs) at 8h and 24h, respectively, after injection of the vaccine (**Fig. 3c, d**). We first interrogated publicly-available databases to delineate putative biological pathways associated with the detected DEGs at 8h and 24h. We found conserved signatures of pathways associated with stress response (e.g, PERK−mediated unfolded protein response or ATF6 activation of chaperone genes), RNA processing (e.g, Insulin−like Growth Factor−2 mRNA Binding Proteins or MAP1K activation), vascular response (e.g., endothelial cells modulation), nervous response (e.g., modulation of axonal guidance), immune response (e.g, positive regulation of type 2 immunity or phagosome activation), and skin homeostasis (e.g, retinoic acid metabolic process or homotypic cell-cell adhesion). Interestingly, at 8h after vaccine injection, we observed the early expression of genes involved in the detection and response to xenobiotic agents, i.e., molecules that are not normally present in animal’s life (**Fig. 3e** **and Table S2**).

We next quantitatively analyzed the expression of genes associated with each biological pathway (**Fig. 3f**). At both 8h and 24h, we found that most DEGs were associated with the biology of keratinocytes or other structural cells. The most upregulated DEGs (25 at 8h and 52 at 24h) were reported to be involved in the global regulation of skin homeostasis such as Late Cornified Envelope 2C^33^ and A (*LCE2C* and *LCE2A*)^34^, pepiplakin-1 (*EPPK1*), the protease inhibitor Serpin Family B Member 13 (*SERPINB13*)^35^, Aspartate Beta-Hydroxylase Domain Containing 2 (*ASPHD2*), Lanosterol Synthase (*LSS*) and Arachidonic acid 12-lipoxygenase (*ALOX12*). A large proportion of the other DEGs was associated with the activation of the immune response in keratinocytes, RNA processing, vascular response, and stress in keratinocytes such as thermo-sensitive transient receptor potential M4 (*TRPM4*)^36^, Cleavage and polyadenylation specificity factor subunit 1 (*CPSF1*)^37^, Metallothionein-1G (*MT1G*) and Calmodulin-like protein 5 (*CALML5*)^38^, respectively. Interestingly, upon injection of the COVID-19 vaccine, we found a global downregulation of IL-6 signaling (i.e., *Il6, IL6R*) and a conserved strong expression of genes associated with the transport of lipoprotein, such as low-density lipoprotein receptor (*LDLR*) and low-density lipoprotein receptor-related protein 8 (*LRP8*), two pathways previously suggested to be involved in LNP particle incorporation^39^. The genes Adrenomedulin-2 (*ADM2*), that encodes a member of the calcitonin gene-related peptide (CGRP)/calcitonin family of hormones, and Protein Kinase C Zeta (*PRKCZ*) were also strongly upregulated and previously reported to be associated with skin inflammatory processes. Finally, we also found a global signature of *MYC*, a key gene that participates in many cellular functions, including cell cycle, survival, protein synthesis, cell adhesion, and microRNA expression^40–42^ (**Fig. 3f**).

Taken together, these data demonstrate that after s.c. injection of the COVID-19 vaccine in natural human skin modules, a global modulation of the human skin ecosystem can be detected at the transcriptomic level. By assessing subtle alterations in gene expression, we can discern the precise sequential events occurring between the exogenous substance and the skin’s biological processes. These events encompass the recognition and response to a foreign agent, the adjustment of skin homeostasis, the activation of stress responses, and the initiation of immune reactions. Thus, by combining skin modules, secretome analysis, and bulk RNAseq analyses, we could gain valuable insights into how natural human skin globally adapts and responds to external stimuli, shedding light on the complex mechanisms governing its physiological and protective functions.

### Single-cell profiling of skin-resident immune cells after subcutaneous vaccination enables the tracking of vaccine sequences and associated modulation of transcriptomic profiles

To gain insight into precise innate mechanisms occurring at the site of vaccine injection, we performed a scRNAseq analysis on dissociated human skin modules enriched in skin-resident CD45^+^ immune cells at 8h and 24h post s.c. vaccine injection (**Fig. S1, lower panel**). Using the Uniform Manifold Approximation and Projection (UMAP) approach, we analyzed the expression patterns of 26,248 genes in 15,564 single cells from the aggregated dataset of the two time points and could unambiguously identify the presence of 14 global immune cell clusters (**Fig. 4a**). All populations were annotated based on their combined expression of cardinal marker genes as previously described^43–45^, such as *CD3E*, *CD4*, *FOXP3*, *CD8A*, *GZMB*, or *KLRD1 for lymphoid cells*, and *ITGAX*, *CD1A*, *CD207*, *TPSAB1*, or *CD68 for myeloid cells* (**Fig. 4b**). We next developed an analytical pipeline, carefully adapted to the amount of biological material contained in each natural human skin module, to sequentially track and quantify the presence of SARS-CoV-2 spike mRNA copies into cells and investigate associated modulations of transcriptomic states at the single cell level (**Fig. S4**).

As the SARS-CoV-2 spike sequence is not naturally present in the 24 chromosomes of the human genome reference database (GRCh38)^46^, we built an “*in-silico* 25^th^ chromosome” at the end of the GRCh38 that corresponds to the sequence of the vaccine (**Fig. S4**). To this end, we created an entry into the GRCh38-associated GTF annotation files and added the entire spike sequence as CDS (CoDing Sequence) and an exonic entry from positions 58 to 3879 to suppress the putative UTR (UnTranslated Region) defined by NAanalytic (the detailed protocol is described in the Methods section). We then aligned the scRNAseq raw data to the custom reference transcriptome, and proceeded to perform conventional analyses. We found that 8h after s.c. injection of the vaccine, the DC/macro cluster, and more surprisingly the mast cell cluster, expressed significant levels of spike mRNA (**Fig. 4c-e**). These data strongly suggest that, in our experimental conditions, the LNPs that vehicle the spike mRNA sequences will be preferentially incorporated into DC/macrophage and mast cell compartments after s.c. injection.

To evaluate the potential impact of s.c. vaccination in different immune populations, we used the *R* package *Augur*, a machine-learning-based algorithm specifically-designed to predict the populations that show the highest cellular response to perturbation between two treatment conditions (in our case, water versus vaccine)^47^. After 8 hours, the *Augur* analysis revealed significant transcriptomic changes in immune compartments, including DC/macrophages, mast cells (both having detectable levels of spike mRNA), LC1s, LC2s, and CD8 T cells (these last populations having very low/undetectable levels of spike mRNA), while at 24h, only the DC/macrophage compartment exhibited continued statistical differences (**Fig. 4f**). We next extracted the list of DEGs associated with our analysis and inferred putative biological pathways regulated in each immune compartment at 8h in DC/macrophages, mast cells, LC1s, LC2s, and CD8 T cells and in DC/macrophages at 24h. Across most of the cell types examined, we found conserved signatures of pathways associated with mRNA processing and translation machinery, type I immune response, Roundabout (Robo) receptors signaling (i.e., involved in actin cytoskeleton remodeling), SARS-CoV-1/2 antiviral response, and cellular stress (**Fig. 4g**). Interestingly, DC/macrophage and LC2 subsets shared some common additional signatures of genes associated with clathrin-mediated endocytosis (i.e., reported to be involved in LNP incorporation), RHO GTPase signaling, MAPK signaling, transcriptional regulation and protein modification, insulin receptor pathways, cell migration, modification of cellular homeostasis, TGF-β signaling, regulation of neuronal compartment and activation of blood/platelet machinery. However, some pathways were found specifically upregulated in the DC/macro subset upon s.c. vaccination, such as FLT3L/FLT3 axis, NF-κB signaling, MHC-I-mediated cross-presentation^48–50^, epidermal growth factor (EGF)/EGF receptor signaling, response to interferon and reactive oxygen species (ROS) production. The LC2 subset also expressed a specific signature of genes associated with FcγR-mediated phagocytosis, MHC-II-mediated antigen presentation and VEGF/VEGFR, and hormone signaling (**Fig. 4g**).

Taken together these data demonstrate that upon s.c. injection of the vaccine, LNP-loaded spike mRNA sequences were detected at 8h, but not 24h following injection, mainly in two innate immune clusters harboring DC/macro and mast cell signatures. Nonetheless, 8h after s.c. injection of the vaccine, we could detect a general modulation of the transcriptomic programs of skin-resident DC/macrophages, mast cells, LC1s, LC2s, and CD8 T cells that was still maintained after 24h in DC/macrophages. In particular, DC/macrophages, and to a lesser extent LC2s, exhibited enhanced expression of genes associated with clathrin-mediated endocytosis, antiviral response, mRNA processing, antigen presentation, and general cellular activation.

**Fig. 4.**
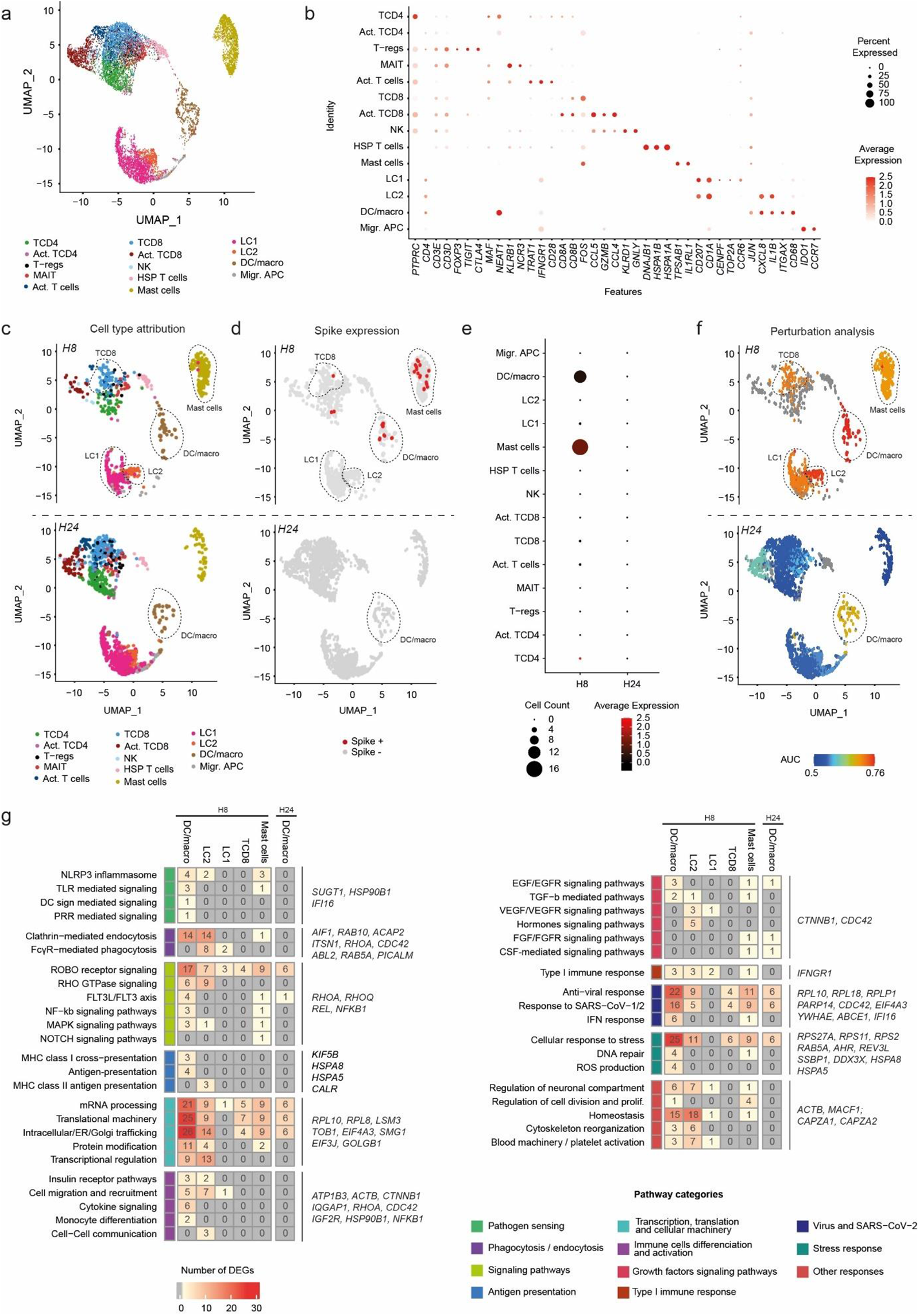
Identification of skin-resident immune cells involved in internalization and response to s.c. injection of COVID-19 vaccine using scRNAseq. **a**, UMAP plots of CD45^+^ skin-resident immune cells in natural human skin modules. Data obtained at 8h and 24h for water and COVID-19 vaccine s.c. injected models were aggregated on the UMAP. Populations were annotated based on the expression of common specific genes. **b**, Dot plot of expression values of cell-type-specific marker genes. **c**, UMAP plots of cell attribution at 8h (top) and 24h (bottom) after injection. **d**, UMAP plots indicating spike RNA detection in each cluster. **e**, Number of cells by cell type which have incorporated spike RNA and levels of expression. **f**, Analysis of gene expression perturbation due to COVID-19 vaccine s.c. injection with *Augur*. The AUC allows the evaluation of levels of gene expression perturbation by cell type (AUC=0.5 no perturbation between conditions, the closer the AUC is to 1, the greater is the difference in gene expression within a cell type between the two conditions). **g**, Biological pathway analysis based on DEGs in the COVID-19 vaccine condition compared to water. *Act. TCD4: Activated CD4 T cells; Act. TCD8: Activated CD8 T cells; Act. T cells: Activated T cells; TCD4: CD4 T cells; TCD8: CD8 T cells; DC/macro: Dendritic Cells/macrophages; HSP T cells: T cells expressing Heat Shock Proteins; LC1: Langerhans Cells type 1; LC2: Langerhans Cells type 2; MAIT: Mucosal-Associated Invariant T cells; Migr. APC: Migratory Antigen Presenting Cells; NK: Natural Killer cells; T-regs: regulatory T cells*.

### Subcutaneous and intradermal deliveries of the vaccine foster the development of mature antigen presenting cells and cell-cell interactions

The spatial organization of the skin immune system across various anatomical layers is well-established. Prior research has indicated that diverse routes of vaccine administration may selectively target specific immune subsets, leading to distinct cellular and humoral immune responses^51,52^. We generated several human skin modules from the same donor as in **Fig. 4** to investigate potential changes in immune cells transcriptomic profiles and incorporation of spike sequences when the mRNA-1273 vaccine was administered i.d. versus s.c. Again, we detected the presence of significant levels of the spike sequence mostly at 8h, but less at 24h, after i.d. vaccine injection in DC/macro and mast cell clusters (**Fig. S5a-c**), confirming a natural capacity for these cells to uptake/be targeted by the LNP-loaded mRNA particles. We next used *Augur* to investigate potential transcriptomic changes in the immune compartment. We found that 8h after i.d. vaccination, DC/macrophages, LC1s, and CD8 T cells, but not mast cells nor LC2s, exhibited significant transcriptomic changes (**Fig. S5d**). However, compared to results observed upon s.c. administration, all of the myeloid compartment, including LC2, migratory APC and mast cell clusters, and a large proportion of CD4 and CD8 T cells underwent significant transcriptomic changes 24h after i.d. administration of the vaccine.

We next applied the algorithm CellPhoneDB to explore differences in cell-cell communication and ligand-receptor interactions between different immune cells when the vaccine was injected s.c. or i.d.^26^. 8h after s.c. injection of the vaccine, we found that myeloid cells, including DC/macrophages, LC1s, LC2s, and migratory APCs exhibited an increased capacity of interaction with other immune cell types within the tissue, including CD4 and CD8 T cell subsets, as compared to the clinical grade water control (**Fig. 5a**). Such a pattern of interaction was then significantly decreased at 24h. Conversely, 8h after i.d. injection of the vaccine, the pattern of interaction of myeloid cells with other immune cell subsets was less pronounced than that observed after s.c. injection. However, 24h after i.d. injection most myeloid cells exhibited a significant increase in their capacity to form interactions with lymphoid cells such as CD4 and CD8 T cell subsets (**Fig. S6a**). We next inferred the predicted ligand-receptor pairs that could be engaged between the different interacting immune cells. We found that most of the predicted interactions were either between myeloid cells or between myeloid cells and CD4 T cells, independent of the route of administration (**Fig. 5b** **and S6b**). Interestingly, we found significantly more predicted pairs of ligand-receptor after s.c. injection than after i.d. injection of the vaccine. Among the predicted interactions, we observed that the s.c. vaccination could favor interactions between CD4 T cells and antigen presenting cells (i.e., clusters DC/macro, LC1, LC2, migratory APCs). Notably, the interactions between CD4 T cells and DC/macrophages specific to vaccine condition were: CD44 (i.e., involved in cell-cell adhesion) and TYROBP (i.e., protein tyrosine kinase involved in cell activation), TNFSF13B (i.e., BAFF involved in lymphocytes activation) and TFRC (i.e., transferrin receptor involved in T cell activation), PTPRC (i.e., CD45) and MRC1 (i.e., mannose receptor C type 1 or CD206), LGALS9 (i.e., Galectine 9) & CD44, IL1B and IL1 receptor (and IL1 receptor inhibitor), IL1B and ADRB2 (i.e., adrenergic receptor beta 2 involved in immune modulation), HGF (i.e., Hepatocyte Growth Factor) and CD44, CXCL8 (i.e., interleukine 8) and NR3C1 (i.e., glucocorticoid receptor), CD55 (i.e., a potent activator of naïve T cells) and ADGRE5 (i.e., CD97 involved in T cell activation), CCL3 and CCR1 (**Fig. 5b**).

We finally investigated changes in the activation status of skin-resident APCs at the protein level using Multiplex ANnotated Tissue Imaging System (MANTIS^®^)^53^, an interactive analytical system based on multiplexed confocal imaging, that automatically generates a digitized version of the skin immune landscape and enables single-cell quantitative data visualization. Using MANTIS^®^, we generated an attribution matrix to automatically annotate dermal DCs (dDCs, CD45^+^ CD1c^+^ CD207^-^), Langerin^+^ dDCs (CD45^+^ CD1c^+^ CD207^+^) and Langerhans cells (LCs, CD45^+^ CD1c^-^ CD207^+^)^54^ (**Fig. 5c,d**) and analyzed the expression of well-established activation/maturation markers such as HLA-DR, CD80, CD86, CD83, CD40, and CCR7 in modules generated from 4 donors and injected s.c. with the vaccine or water for injection control. A total of 8,166 single CD45^+^ immune cells were identified with the following distribution 7.4% dDCs, 26,7% Langerin^+^ dDCs, 33,9% LCs and 32% other immune cells across the donors (**Fig. 5e, f**). We found that LCs and Langerin^+^ dDCs significantly upregulated most of the maturation markers mentioned above at 24h, but not 8h following vaccination (**Fig. 5g** **and S7**). We also observed that LCs, that are classically detected in the epidermis, mostly relocated to the dermal area 24h after vaccination (**Fig. 5f**). However, dDCs seemed to be less responsive to the vaccine and upregulated CD83 and CCR7 after 24h.

Taken together, these results provide robust evidence that the mRNA-1273 COVID-19 vaccine maintains a relatively consistent tropism for dermal DC/macrophages and mast cells when analyzed by scRNAseq, regardless of the route of administration in natural human skin modules. These data also suggest that skin-resident myeloid cells acquire a mature phenotype of APCs after vaccination (in particular LCs and Langerin^+^ dDCs), while increasing their predicted interactions with surrounding CD4^+^ T cells with specific pairs of ligands and receptors (**Fig. 6**). Here we show that the VaxSkin pipeline is uniquely positioned to assess numerous scientific questions in the field of human innate immune response to vaccines, not only in DCs, but also in a wider range of native tissue-resident myeloid and lymphoid cell populations depending on the route of administration.

**Fig 5.**
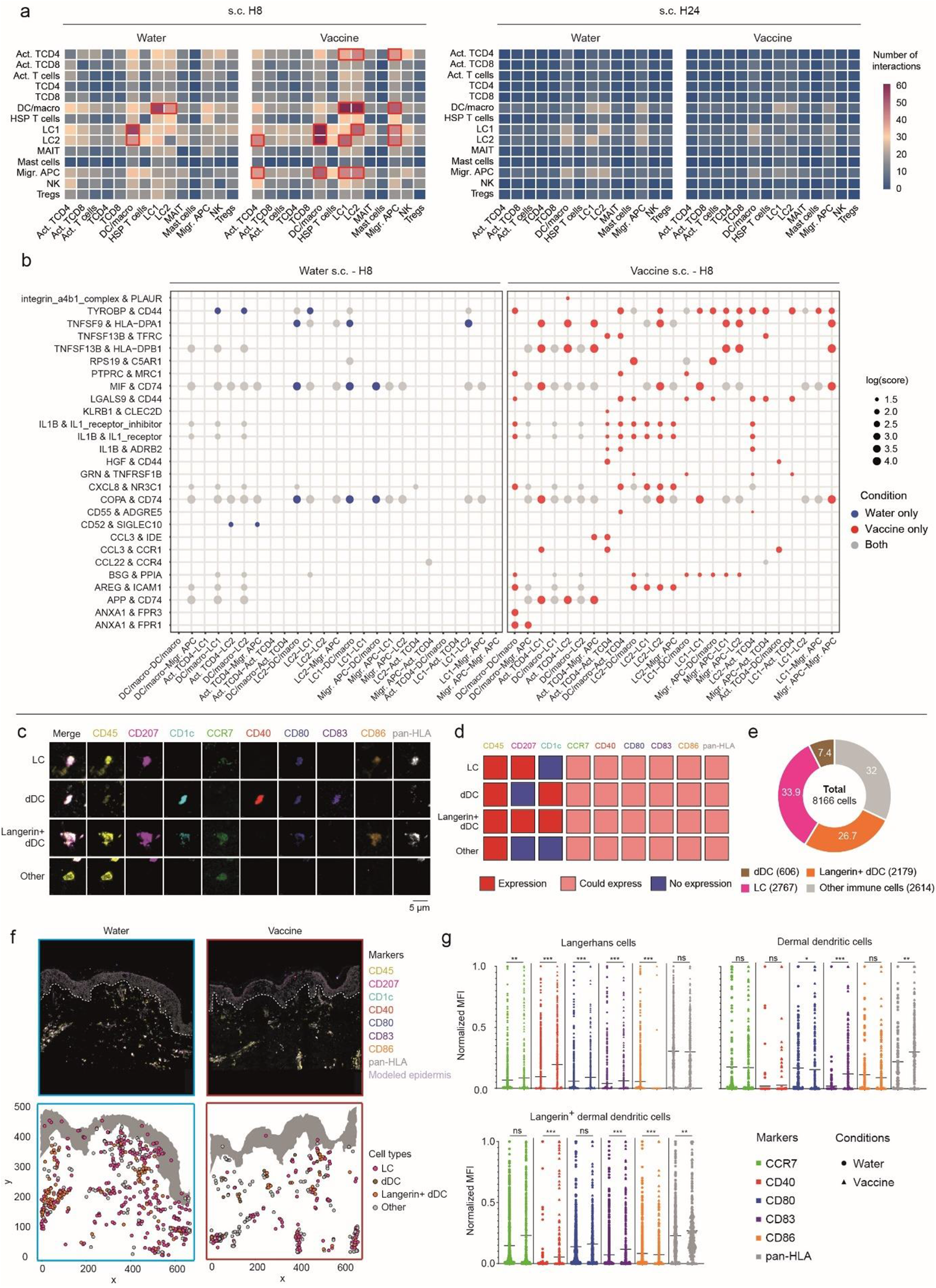
Modifications in interactome signatures emerged 8h post-s.c. injection of the COVID-19 vaccine, preceding APCs activation manifestation within 24h. **a**, Analysis of the number of interactions between immune cell types, 8h and 24h after s.c. injection of water or COVID-19 vaccine. Red squares indicate a number of interactions superior to 35. **b**, Dot plot of the ligand-receptor interaction scores between immune cell types most affected by vaccine injection (i.e., for which the absolute value of number of interactions between vaccine and water were above 10) 8h after s.c. injection. Only statistically significant scores are presented. **c**, Examples of single-cell staining on human skin cryosection used for APCs attribution and activation panel. Scale bar, 5 μm. **d**, MANTIS^®^ attribution matrix for APC activation panel. **e**, Total number of immune cells analyzed on skin sections and proportions of analyzed LC, dDCs, and Langerin+ dDCs. **f**, Representative 3D confocal multiplex images of 40µm thick skin cryosections (top) and associated digital maps (bottom) of selected markers of the MANTIS^®^ APC activation panel of s.c. water-(left) and vaccine-injected (right) human skin modules, 24 hours post-injection. **g**, Dot plots of APC activation markers (CCR7, CD40, CD80, CD83, CD86, and pan-HLA) normalized mean fluorescence intensity (MFI) in LCs (top left), dDCs (top right) and Langerin^+^ dDCs (bottom left) in water-(left) and vaccine-injected (right) human skin modules, 24h post-injection. *Act. TCD4: Activated CD4 T cells; Act. TCD8: Activated CD8 T cells; Act. T cells: Activated T cells; TCD4: CD4 T cells; TCD8: CD8 T cells; DC/macro: Dendritic Cells/macrophages; HSP T cells; T cells expressing Heat Shock Proteins; LC1: Langerhans Cells type 1; LC2: Langerhans Cells type 2; MAIT: Mucosal-Associated Invariant T cells; Migr. APC: Migratory Antigen Presenting Cells; NK: Natural Killer cells; T-regs: regulatory T cells; LC: Langerhans cell; dDC: dermal dendritic cell*.

## Discussion

Here we introduce the VaxSkin platform, a versatile framework primarily designed for longitudinal profiling of the human skin ecosystem in response to vaccines at the injection site. This platform comprises of a chemically-defined gel matrix, combined with a topical silicone ring, ensuring skin viability and immunocompetence for at least 7 days. Each individual human skin module consists of the matrix-skin biopsy-silicone ring association, allowing s.c. or i.d. delivery using standard clinical syringes while preserving the integrity of native adipose tissue. Compared to classical skin-related data obtained during clinical trials in humans, this method offers an remarkable scalability, enabling multiple parallel experiments in one donor and extension to cohorts based on selectable human biological diversity. The platform incorporates an analytical pipeline that is tailored to the amount of biological material found in a single module. It utilizes integrated multiparametric analyses to extract both immune and structural datasets from each module. Additionally, it employs an *in silico* approach to selectively track and quantify mRNA vaccine incorporation at the single-cell level.

Using the VaxSkin platform, we characterized the biological response of the skin, at both organ and single-cell levels, in response to injection of the mRNA-1273 COVID-19 vaccine. At the organ level, our data strongly support the notion that injection of the vaccine induces comprehensive modulation of the human skin ecosystem at the secretomic and transcriptomic levels. Through the analysis of gene expression levels over time, we could discern the precise sequential events that unfold between the exogenous substance and the skin biology. These events encompass the recognition and response to the foreign agent, adjustments in skin homeostasis, activation of stress responses, and initiation of immune reactions. It is very interesting to speculate that the injection of a vaccine into the skin not only affects APCs, but also generates a complex biological response at the organ level that is likely to influence the efficacy of the immune response to carry out protection (**Fig. 6, left**). Exploring and manipulating the mechanisms that govern the global regulation of the skin ecosystem following vaccination, extending beyond the targeted immune cells, holds great promise for future investigations.

When analyzed through the lens of immunology at the single-cell level, we found that, under our experimental conditions, the vaccine consistently targets dermal DC/macrophages and mast cells, irrespective of the administration route in natural human skin modules (**Fig. 6, right**). One set of genes expressed by the DC/macrophage subset after vaccination was linked to clathrin-mediated endocytosis, which is one the pathways reported to be involved in LNP internalization^55,56^. However, this set of genes was not found to be upregulated in mast cells, despite their high propensity to incorporate vaccine LNP. Mast cells are highly granular/vesicular cells^57–59^, it would be interesting to gain a better understanding of how they could incorporate such quantities of LNP. These findings are in line with previous reports that LNP could efficiently transfect *in vitro*-cultured mast cells^60,61^. While DC/macrophages are professional APCs that can initiate the activation of the adaptive immune response, less is known about the capacity of mast cells to present antigens to T cells in the context of vaccination. Previous work suggests that mast cells could activate antigen-experienced CD4 T cells in mice and humans^62–64^, to which extent such findings are applicable in the context of vaccination is still unknown. The primary function of mast cells is to release granule-associated mediators, a process named “degranulation”^65^ and involved in drug-associated injection site reactions^66,67^. The COVID-19 vaccine has been reported to induce redness, swelling, and pain in the skin after injection^68^, the extent to which these clinical manifestations are directly associated with the capacity of mast cells to incorporate LNPs remains an open question.

Beyond the above-described spike-positive immune cells, the infusion of the vaccine also triggered substantial transcriptomic changes in the entire myeloid compartment, including LC2, migratory APC, mast cell clusters, and a significant proportion of CD4 and CD8 T cells. It is possible that our *in silico* detection of the spike is not sensitive enough to catch smaller incorporation of the vaccine in those cell types. However, in line with our organ level analysis, it is likely that the vaccine triggers a qualitative modification of the skin ecosystem and fine tuning of “bystander non-targeted cell types” at the site of injection. It is interesting to note that CD8 T cells were the most affected lymphoid cell type and, currently, the investigation of how tissue-resident CD8 T cells sustain themselves in the absence of antigen presentation has become a rapidly advancing field in T cell immunology^69^.

Finally, previous reports in the field have shown that vaccine efficacy could be affected by the route of administration^51,70,71^. Using the VaxSkin platform, we found that the i.d. injection route shows a prolonged modulation of both myeloid and lymphoid cells compared to the s.c. route, persisting from 8h to 24h after vaccine administration. These findings further demonstrate that the route of administration could lead to diverse effects on local immune cell populations within the skin, involving not only dermal DC/macrophages, but also a broader array of myeloid and lymphoid cells. Gaining deeper insights into how the route of injection influences immune cell targeting and the potential for fine-tuning the overall regulation of the skin ecosystem after vaccination in humans holds significant promise for future research. Such multi-modal vision of skin vaccination could pave the way for the development of more effective and tailored vaccine development strategies with improved immune responses.

There is a critical need for scalable technological platforms to evaluate the local immunogenicity and safety of drug and vaccine candidates in a setting that faithfully replicates a fully human environment before advancing to clinical trials. Here we show that the use of the natural human skin module technology, coupled to a scalable analytical pipeline, has great potential to advance the study of skin physiology. In the context of drug and vaccine discovery and development, it will be especially valuable for the study of molecular mechanisms of action, prioritization of lead candidates, toxicity and safety testing, and biomarkers identification.

**Figure 6.**
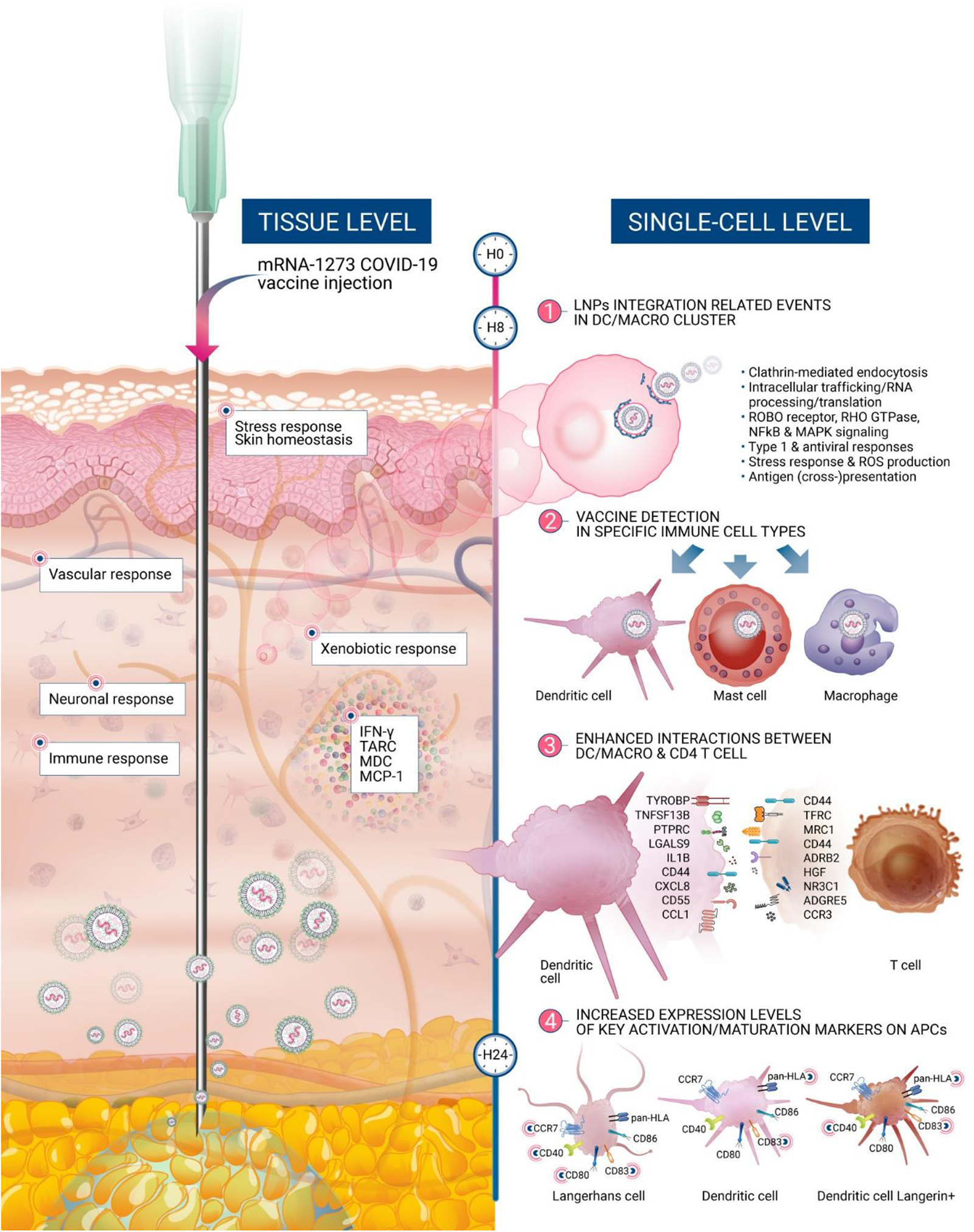
VaxSkin solution provides details on the mechanism of action of mRNA-1273 COVID-19 vaccine in human skin. The ability to inject the vaccine in immunocompetent human skin module coupled with a multi-omics approach, which combined multiplex cytokine measurements, multiplex imaging, bulk and single cell RNA seq, enables data integration at the tissue and single cell levels.

## Methods

### Natural human skin modules: sourcing, production, treatment, and sampling

Skin biopsies were obtained from Genoskin SAS (https://www.genoskin.com/). Genoskin has obtained all legal authorizations necessary from the French Ministry of Higher Education, Research and Innovation (AC-2017-2897) and the Personal Protection Committee (2017-A01041-52). Natural human skin modules of 23mm with 10 mm total thickness and 18/23 mm diameter silicon rings were embedded in a proprietary solid matrix, with the epidermal surface in direct contact with ambient air. The natural human skin modules were mounted on cell culture inserts, loaded in multi-well companion plates with the lid containing 3mL of chemically-defined xeno-free Genoskin medium. The natural human skin modules were cultured at 37°C in a standard incubator (5% CO2) up to 10 days, and culture medium was renewed each day.

#### Modules viability study

At day 0, 3, 5 and 10, the modules were sampled and saved either in OCT for immunostaining, or in FFPE for H&E staining, or processed for single cell RNA sequencing (**Fig. S1, upper panel)**.

#### VaxSkin study

At 0h, natural human skin modules were injected subcutaneously either with clinical-grade water or mRNA-1273 COVID-19 vaccine (Moderna). Two sets of experiments were performed. Both gather non-injected, water-injected or vaccine-injected conditions. The first set of experiment was sampled at 0, 8 and 24h, and split into 4 pieces: 1⁄4 of the module (including the hypodermis) was saved for paraffin embedding, 1⁄4 of the module without hypodermis was frozen in RNA latter and stored at – 80°C, 1⁄4 of the module without hypodermis was embedded in OCT and stored at -80°C, and 1⁄4 of the module without hypodermis was frozen and stored at -80°C. The matrix and the culture medium of the module was collected and stored at -80°C. The second set of experiment was collected at 0, 8 and 24h and processed for single cell RNA sequencing. Two models were used for each condition (**Fig. S1, lower panel)**.

### Histology

5 µm formalin-fixed and paraffin embedded (FFPE)-tissue sections were rehydrated by immersing the slides in the following solutions: two washes of xylene (4 minutes each), two washes of 100% ethanol (2 minutes each), one wash of 95% ethanol (1 minute) and three washes of PBS. The slides were then stained with hematoxylin & eosin and dehydrated by immersion in 99%-100% ethanol solution and xylene. The slides were covered with mounting medium (Eukitt^®^, Orsatec), and sealed with a coverslip. For each sample, representative images were acquired at 40x magnification with a DMi1 (Leica) equipped with a MC170 HD camera (Leica).

### Injection and bolus observation

The natural human skin modules were injected subcutaneously or intradermally with 100μL of patent blue V (E131), 0.5% (w/v) in PBS, or KI solution just before the picture acquisition or tomography acquisition respectively. For the tomography acquisition, an EasyTom X-ray microtomography machine (RX Solutions) was used. This machine is equipped with a 150 kV microfocus X-ray source (Hamamatsu). X-ray radiographs were taken with a beam power of 14 W, corresponding to an acceleration voltage of 65 kV and a current of 220 µA. The exposure time was set to 0.15 s. X-ray tomographies were obtained by acquiring 1440 images at different angles evenly distributed over 360°. The 3D volume was reconstructed using RX Solution Software (X-Act). Post-processing of the 3D images was performed using Imaris software. The diffusion area of the contrast agent, the hypodermis and the epidermis/dermis were manually segmented using 2D views, before performing 3D volumes reconstruction.

### Thick section staining and biphoton imaging

100µm cryo-sections were fixed with 4% PFA for 30min. Slides were rinsed, blocked and permeabilized with PBS 0.5 % (w/v) BSA (Sigma-Aldrich), 0.3 % Triton X-100 (Merck) for 1 hour at room temperature, then incubated with fluorophore-coupled antibodies or unconjugated antibodies for 24h at 4°C in the dark. The slides were rinsed with PBS 0.5 % (w/v) BSA, 0.3 % Triton X-100 and incubated with the secondary antibodies for 2h at room temperature in the dark. Finally, the slides were rinsed and mounted in Mowiol medium (Sigma-Aldrich) and sealed with a coverslip. All conjugated and unconjugated antibodies used in this study were validated in single immunostainings (unpublished), and are listed in **Table S3**. Mosaic images were acquired using an Ultima 2P plus biphotonic microscope (Bruker Corporation) equipped with a laser Chameleon Discovery NX (Coherent Inc) operating at a 770nm excitation wavelength, and a 20x objective, 1 Numerical Aperture, 2mm WD (Olympus,). The acquired images were stitched and processed using the Imaris Software.

### MSD analysis

Levels of cytokines secreted in the matrix were quantified using the 36-plex kit from Meso Scale Discovery (MSD), following manufacturer’s instructions. Cytokine concentrations expressed in pg/mL were calculated by comparison to the standard curve which was generated in the same biological matrix as the samples. Only values that were in the detection range were taken into account. The statistical analysis was performed using an in-house application. For each detected cytokine, Shapiro-Wilk and Fisher tests were used to assess the normality and homogeneity of the data, respectively. A Student t-test, Mann-Whitney-Wilcoxon or Welch’s t-test (depending on preliminary statistical tests) was then performed to evaluate the significance of the mean’s differences. Cytokines were then selected according to significant changes over the 6 analyzed donors. This selection was used as an input to *PathfindR*^72^, version 2.1.0, R package with default parameters, to perform an active subnetwork enrichment analysis approach that represents enriched pathways. Pathways databases Gene-Ontology^73^ biological process, KEGG^74^ and Reactome^75^ and Protein Interaction Network Biogrid and String were used to compute the enrichment.

### Bulk RNA sequencing

Briefly, a 4mm punch of skin was digested in Trizol using a GentleMACS™ Octo Dissociator (Miltenyi Biotec) to extract RNA from the tissue. The RNA was then purified using a RNAeasy Plus Mini Kit (Qiagen) following the manufacturer’s guidelines. The concentration and quality of RNA were determined in each sample using a NanoDrop (ThermoFisher Scientific). The extracted RNA was stored at -80°C until library preparation. mRNA library preparation was realized following manufacturer’s recommendations (Illumina Stranded mRNA Prep from Illumina). Final samples pooled library prep were sequenced on Novaseq 6000 Illumina with SP-200cycles cartridge (2x800Millions of 100 bases reads), corresponding to 2x30Millions of reads per sample after demultiplexing. Read quality control was performed with *MultiQC*^76^, version 1. All samples passed this quality check, and reads were aligned with *STAR*^77^ version 2.7.2b on the human GRCh38 reference genome. Aligned reads were ordered and indexed using SAMtools^78^, version 1.16.1. Genes counts were computed with HTSeq^79^ counts version 0.12.4. *DESeq2* version 1.40.2 was used to process raw count. *DESeq* object was created from a sample table with a design that takes into account the condition (treatment and time point) and the donor. A variance stabilization was performed on gene count using the vst function, to plot samples on PCA. Donor and gender biases were corrected using *Combat-seq*^31^ from R library *sva*^32^ version 3.48 with default settings. Corrected count matrices were analyzed with *DESeq2* version 1.40.2. Genes with an adjusted p-value (Benjamini-Hochberg correction) lower than 0.1, a log2 fold change greater than 0.26 and lower than -0.26 were considered as significant. These genes were used as an input to *PathfindR* R package, version 2.1.0, with default parameters to perform an active subnetwork enrichment analysis approach that represents enriched pathways. Molecular function and biological process of Gene-Ontology, KEGG and Reactome pathway databases were used to compute the enrichment.

### Single cell RNA sequencing

Briefly, skin was harvested in a pre-digestion medium, fragmented into pieces and incubated at 37°C on a rotating plate to remove impurities. The supernatant was discarded and the samples were digested for 30 minutes on a rotating plate with 1.25 mg of Liberase (Sigma Aldrich) and 2.5 mg of DNAse I (Sigma Aldrich) to disaggregate the tissue. Samples were further dissociated with the GentleMACS™ Octo Dissociator. Cells were then enriched with a Percoll gradient and washed. For the *modules viability study*, the cell suspension resulting from the dissociation was sorted using a FACSymphony (BD Biosciences) cytometer to isolate Sytox^-^negative alive cells. For *VaxSkin study*, the cell suspension resulting from the dissociation was enriched in CD45^+^ cells with an EasySep Release Human CD45 Positive Selection Kit (Stemcell Technologies) following the manufacturer’s guidelines. For scRNA-sequencing, cells were encapsulated into droplets using Chromium Next GEM Single Cell 3′ Reagent Kits v3.1 with single indexing, according to manufacturer’s protocol (10× Genomics). Briefly, after generation of nanoliter-scale Gel bead-in-EMulsions (GEMs) using Next GEM Chip G, GEMs were reverse-transcribed in a C1000 Touch Thermal Cycler (BioRad) programed at 53°C for 45 min, 85°C for 5 min, and held at 4°C. After reverse transcription, single-cell droplets were broken and cDNA was isolated and cleaned with Cleanup Mix containing DynaBeads (Thermo Fisher Scientific). cDNA was then amplified with a C1000 Touch Thermal Cycler programed at 98°C for 3 min, 12 cycles of (98°C for 15 s, 63°C for 20 s, 72°C for 1 min), 72°C for 1 min, and held at 4°C. Subsequently, the amplified cDNA was fragmented, end-repaired, A-tailed, index adaptor ligated, and cleaned with cleanup mix containing SPRIselect Reagent Kit (Beckman Coulter) in between steps. Post-ligation product was amplified and indexed with a C1000 Touch Thermal Cycler programed at 98°C for 45 s, 13 cycles of (98°C for 20 s, 54°C for 30 s, 72°C for 20 s), 72°C for 1 min, and held at 4°C. All the libraries were finally cleaned up with SPRIselect beads, and controlled on a Fragment Analyzer HS-NGS run (Agilent). 10x libraries were pooled and charged with 1% PhiX on one SP lane of the NovaSeq 6000 instrument (Illumina) using the NovaSeq 6000 SP Reagent Kit v1.5 Raw sequencing data of the *modules viability study* were processed with *Cellranger*^80^, version 6.1.1, from 10X Genomics. Transcripts were mapped on the reference human genome GRCh38, assigned to individual cells from the specific barcoding and all samples were aggregated. For *VaxSkin study*, water samples were processed similarly to the *modules viability study* data. Regarding the vaccine-injected samples, a custom genome reference was created with *Cellranger mkref* function. In order to detect the exogenic spike COVID-19 mRNA, the sequence retro-engineered by NAalytics^81^ was used and added at the end of the GRCh38 genome as an artificial vaccine ‘chromosome’. An entry into the GRCh38 associated GTF annotation files was created. The entire spike sequence was added as CDS (CoDing Sequence), and an exonic entry from positions 58 to 3879 was also added to suppress the putative UTR (UnTranslated Region) defined by NAanalytic **(Fig. S4)**. Removal of ambient RNA was then performed on *Vaxskin* data with *Cellbender*^82^ version 0.2.2.

The *modules viability* and *VaxSkin* datasets were filtered to remove low-quality or dead cells, by using the *Seurat* R toolkit^83^ version 4.3.0.1. Cells with mitochondrial gene expression higher than 12%, and cells with less than 500 or more than 3100 UMI-counts were removed (**Table S4**).

#### Modules viability study

For each cell, the feature expression measure was normalized by the total expression, multiplied by a scale factor of 10.000 and the result was log-transformed. Feature level scaling was also performed, each feature was centered to have a mean at 0 and scaled by the standard deviation of each feature. To reduce the dimensionality of the dataset, the top 2000 of most variable features were determined from the average expression and dispersion. From these selected genes, Principal Components Analysis (PCA) was performed and the dimensionality of the dataset was determined by using elbow plots that rank principal components by their percentage of variance.

#### VaxSkin study

Normalization and variance stabilization were performed with SCtransform seurat function^84^ using default parameters. Data were integrated using *Harmony*^85^ version 0.0.1 using the top 13 dimensions, four covariate parameters (sample, treatment, timepoint and site of injection) and associated lambda (0.2,1,1 and 1, respectively) were set.

From filtered and preprocessed data, K-Nearest Neighbors (KNN) algorithm was constructed to link cells with similar feature expression and create communities from the Euclidean distance of the PCA space and edge weights refining based on the shared overlap in their local neighborhoods (Jaccard similarity). Cells were then clustered using the Louvain algorithm that optimizes the standard modularity with a resolution value of 0.3 and 1.2 for *VaxSkin* and *modules viability* datasets, respectively. From the most important principal components (15 and 25 for *VaxSkin* and *modules viability* datasets, respectively), t-Distributed Stochastic Neighbor Embedding (t-SNE) were calculated to visualize and explore the dataset. Seurat cell clusters were attributed as cell types by using literature-based specific markers. A sub-clustering on myeloid, lymphoid and structural cells was performed on *VaxSkin* dataset to improve the cell type attribution. Myeloid, lymphoid and structural datasets were integrated using the first 14 principal components for myeloid and lymphoid datasets, and 10 for the structural datasets. Clustering was performed using the same method with a Louvain resolution of 0.3 for all cell types.

Comparisons between days (*modules viability study)* and between water and vaccine conditions (*VaxSkin study*) were performed using a logistic regression model, and the roc method that calculates the classification power of a random forest classifier, respectively. Both datasets were analyzed with the Seurat FindMarkers function. Only genes expressed in at least 10% of cells in each cluster were used to compute the statistical analysis. Genes with an adjusted p-value (Bonferroni correction) lower than 0.05, and, a log2 fold changes greater than 0.58 and lower than -0.58 for the *modules viability* dataset, or a classification power greater than 0.7 and lower than 0.3 for *VaxSkin* dataset, have been considered as significant. These genes were used as an input to *PathfindR* R package version 2.1.0, with default parameters to perform an active subnetwork enrichment analysis approach that represents enriched pathways. Gene-Ontology^73^ molecular function and biological process, KEGG and Reactome pathway databases were used to compute the enrichment.

Pseudotimes of keratinocytes differentiation in the *modules viability study* dataset were calculated with the *Monocle3* R package^86^ version 1.3.1. The pre-processed keratinocytes subset and the most variable genes from Seurat extracted with their annotations were used as an input in *Monocle3*. A matrix readable by monocle was created to build a cds *Monocle3* object from the *new_cell_data_set* function. From this cds, monocle dimensional reduction method and clustering were performed and replaced by their Seurat equivalence in order to use the same dimensional reduction. Trajectory was inferred using default parameters of Monocle after dimensional reduction and cell ordering. *Seurat AggregateExpression* function was used to create two matrices with gene expression from the *modules viability study* dataset: one at the whole tissue level and one at the cell-type level. From these matrices, *Deseq2*^87^, version 1.40.2, was used and only genes with at least 10 counts were conserved. The data were normalized with the rlogTransformation function and a correlation matrix was created and plotted as a correlation heatmap.

Perturbation analysis was performed using the cell type prioritization R library *Augur*^47^, version 1.0.3, with default parameters. By using a random forest classifier, *Augur* assesses the probability of each cell to belong to a specific condition. This prediction was evaluated in cross-validation, and quantified using the Area Under the receiver operating characteristic Curve (AUC).

Interactome study was performed with *CellphoneDB*^26^, version 3.1.0. For each cluster, only genes expressed in at least 25% of the cells of the cluster were conserved. Using its detailed receptor–ligand database, *CellphoneDB* determines the mean interaction value between ligand-receptor complexes between each cell type. Cells are randomly permuted over 1000 iterations, and, at each iteration, the average of the mean expression of receptors and the average of the mean expression of ligand were calculated. The proportion of mean equal or higher than the real mean was calculated and p-values were attributed for each ligand-receptor pair highlighting the specificity of a ligand-receptor complex. If the p-value of enrichment is lower than 0.05, the pair was considered as significant. For each condition, a table gathering all interaction pairs between all cell types was created based on the CellphoneDB ‘significant_means’ output. By substracting control sample to his corresponding vaccine sample, only celltypes with a number of interactions above 10 (in absolute value) were kept, namely: DC/macrophages, activated CD4 T cells, LC1, LC2, and migratory APC. The data were then subjected to a log2 transformation. Dotplots were generated using the *ggplot2* R library version 3.4.3, with the point size representing the log2 of the significant means.

### MANTIS analysis

The MANTIS analysis was performed as in ref ^53^. OCT-tissue sections were blocked and permeabilized with phosphate-buffered saline (PBS), 0.5% (w/v) bovine serum albumin (BSA; Sigma-Aldrich), and 0.3% Triton X-100 (Merck) for 1 hour at room temperature and then incubated with fluorophore-coupled antibodies overnight at 4°C in the dark. The sections were then washed three times in PBS 0.5% (w/v) BSA, 0.3% Triton X-100. Finally, samples were treated with an autofluorescence quenching solution named TrueVIEW (Vector Laboratories) for 5 min. The slides were mounted in Mowiol medium (Sigma-Aldrich) and sealed with a coverslip. All conjugated antibodies used in this study were validated in single immunostainings of human skin and are listed in **Table S5**.

Z-stack images (512 × 512 pixel; 1 μm) were acquired using a confocal microscope SP8 (Leica Microsystems) equipped with an HC PL APO CS2 with 40× numerical aperture 1.3 oil objective, an ultraviolet diode (405 nm), and four lasers in visible range wavelengths (405, 488, 532, 552, and 635 nm). The setup was made up of five detectors [three hybrid detectors with high quantum yield compared to classical photomultiplier (PMT) detectors and two PMTs. Mosaic sequential images were acquired using the between-stack configuration to simultaneously collect individual nine channels and tiles before merging them to obtain one single image. Use of the between-stack configuration and the modulation of the detectors’ detection windows help reduce the leaking of fluorophores.

The mosaic multicolor image was obtained and exported into a .lif format. 3D mosaic images were then imported into Huygens software (SVI) to correct the signal by applying deconvolution and cross-talk correction. The deconvolution was performed using the deconvolution wizard. For each marker, SNR and background signal levels were determined and applied to all images from the same donor. Automatic cross-talk correction estimations were obtained. 3D mosaic images were imported into Imaris software (Oxford Instruments) to separate objects (cells) using a 3D surface segmentation. Before creating the surface objects in Imaris, classical image processing was required. For instance, thresholds were applied to clean the background. Then, segmentation was applied on the CD45 channel surface. For each object, MFI of each marker was extracted. The APCs panel is based on cell type attribution markers (CD45, CD207 and CD1c) and activation markers (CCR7, CD40, CD80, CD83, CD86 and pan-HLA). For bioinformatic analysis, a threshold for each cell type attribution marker was determined to define if a cell is positive or not. For activation markers, the MFI value is raw, to evaluate the level of activation. The epidermis was identified using the natural autofluorescence of the tissue. On the basis of the autofluorescence found in multiple appropriate channel an epidermis surface was created using the Object Creation semi-automated tool of the Imaris software. The coordinates of the epidermis were then exported into .csv format. Statistical properties of each segmented object (cell) in the processed 3D Imaris Multiplex image were automatically calculated. Object area, xyz position, and MFI in all channels were exported as a .csv table.

Cell types were attributed based on the MFI of CD45, CD207 and CD1c. Distinct thresholds were determined for each of these markers and depending on the skin compartment (epidermis and dermis) to allow the removal of autofluorescence signal and background that can be different depending on the compartment (**Table S6**). MFI was corrected by subtracting those specific thresholds and cell types were attributed. For each timepoint, vaccine and water data were statistically compared following MFI transformation. The transformation was made following the formula: (MFI_Marker – min(MFI_Marker)) / (max(MFI_Marker) – min(MFI_Marker))) to represent data between 0 and 1. For each couple vaccine-water samples, preliminary parametric tests of the adequacy to a normal distribution (Shapiro test) and variance homogeneity (Fisher F test) were performed to determine the most relevant statistical test to compare means (Student t-test, Welch’s t-test or Wilcoxon signed-rank test). Statistical tests were performed using Prism 8 (GraphPad Software) and the *Rstats* R package.

## Data visualization

Visualization charts were obtained using the *ggplot2*, *Pigengene*, *pheatmap* and *ComplexHeatmap* R packages, and *matplotlib & seaborn* Python packages.

## Authors’ contribution

N.G. conceived the project. M.S., M.P., E.B., J.M., E.P. and N.G. were involved in experimental design. M.S., M.P., E.B., J.M., E.P. and N.G. performed most experiments and compiled the data. N.S. A.L., A.B. and L.B. provided important help with experiments. E.M. and P.D. provided expertise. All authors participated in analyzing the data and writing or editing the paper. We thank Anthony David, Julie Charpentier, Margot Romero, Anne Dalapa-Amana, Lévi Da Silva, Alexandra Ochando and Hervé Huchon at Genoskin SAS, and Paul Duru and Laurent Malaquin from the LAAS CNRS for discussions and technical assistance. We thank Sophie Allart and Simon Lachambre for technical assistance at the cellular imaging facility of Inserm UMR 1291, Toulouse. We thank Samantha Milia and Timothé Durand-Plavis for technical assistance at the experimental histopathology platform US06/CREFRE. We thank Anne-Laure Iscache, Valerie Duplan-Eche and Fatima-Ezzahra L’Faqihi for technical assistance at the flow cytometry core facility of INSERM UMR 1291, Toulouse. We thank the GeT-Santé facility (I2MC, Inserm, Génome et Transcriptome, GenoToul, Toulouse, France) for the advice and technical contribution to the single cell and bulk RNA sequencing experiments. This work also benefited from equipment and services from the iGenSeq core facility (Genotyping and sequencing), at ICM.

