## Supplementary information for "Multi-modal profiling of biostabilized human skin modules reveals a coordinated ecosystem response to injected mRNA-1273 COVID-19 vaccine"

**Title**

### Supplementary Figures and legends

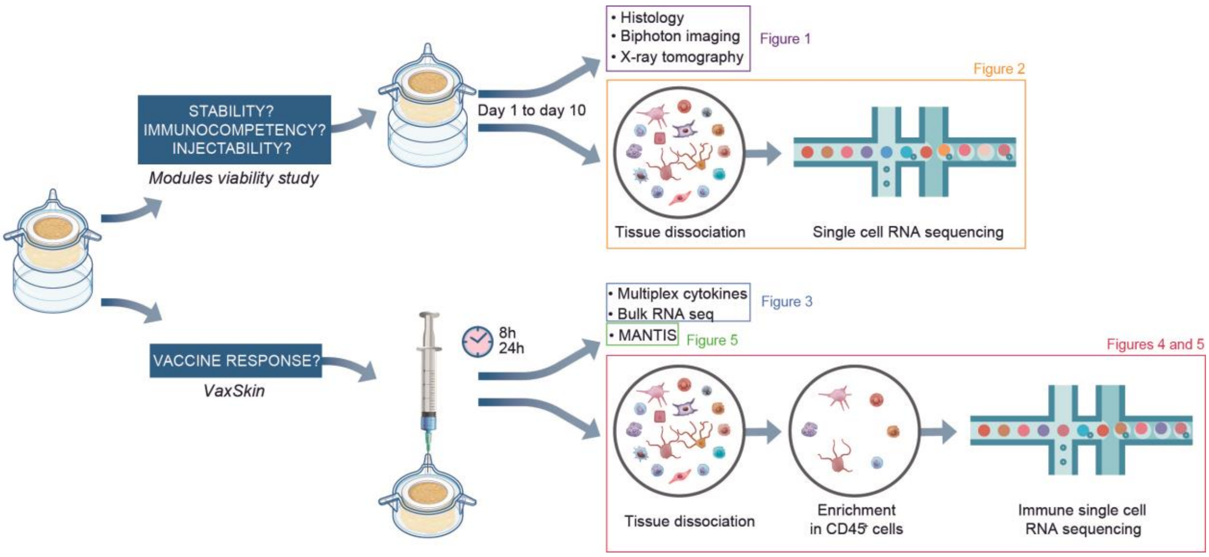


**Figure S1. Analysis pipeline for evaluating stability and vaccine response after mRNA-1273 COVID-19 vaccine injection in natural human skin modules.** The *Modules viability* study involved histology, biphoton imaging, and single-cell RNA sequencing to assess module stability and immunocompetence within 10 days of culture. The immunological reaction to the vaccine was examined using multiplex cytokine measurement and bulk RNA sequencing for an overall immune response perspective. Single-cell analysis techniques, including MANTIS and single-cell RNA sequencing, were employed to investigate the cellular responses triggered at 8- and 24-hours after vaccination.

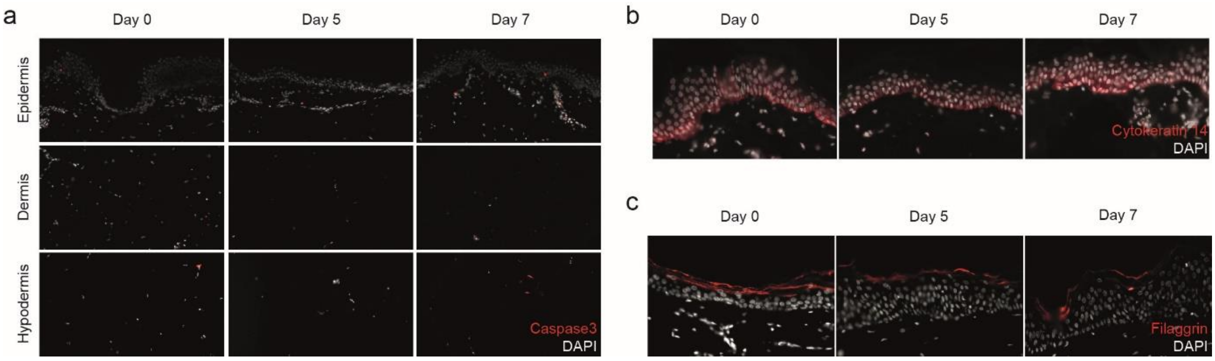


**Figure S2. Natural human skin module's structure and viability assessment over 7 days of culture.** **a**, Immunostaining of DAPI and Active caspase 3 at day 0, 5 and 7 of culture. Three representative images were acquired in the epidermis, dermis, and hypodermis respectively. **b**, Immunostaining of DAPI and cytokeratin 14 at day 0, 5 and 7 of culture. One representative image was acquired in the epidermis. **c**, Immunostaining of DAPI and filaggrin at day 0, 5 and 7 of culture. One representative image was acquired in the epidermis.

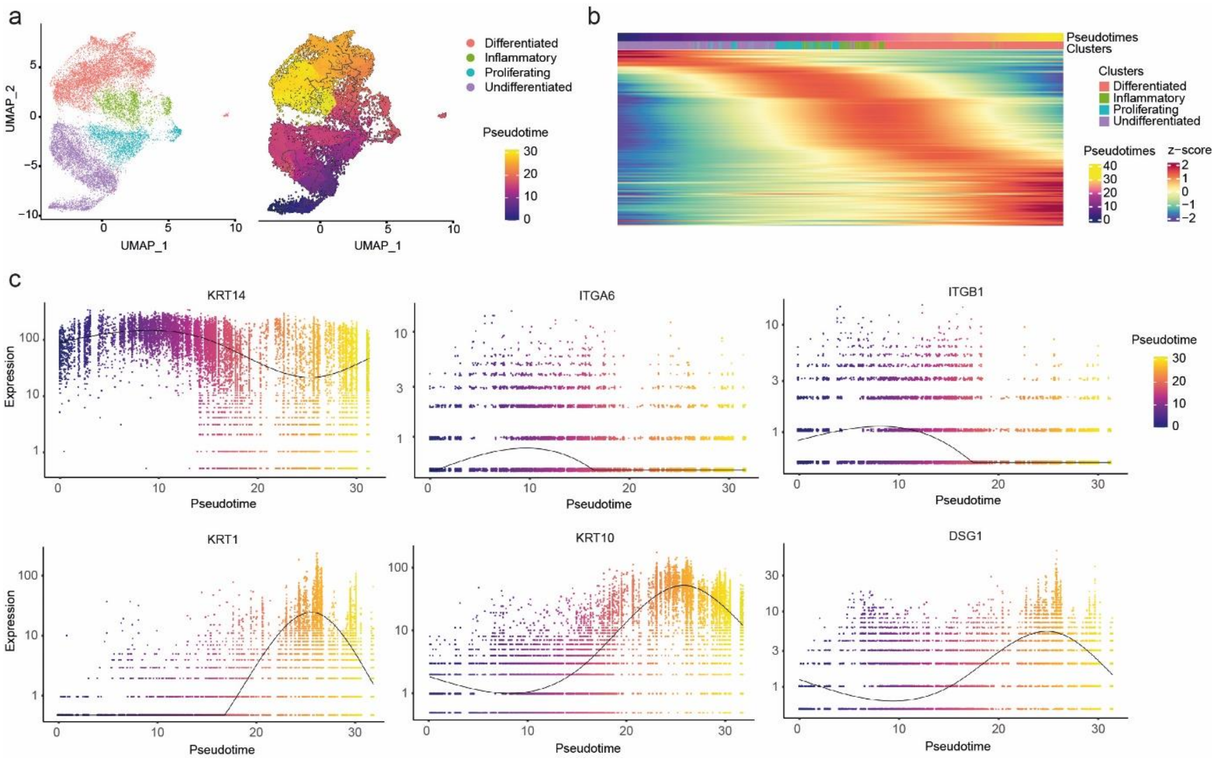


**Figure S3. Pseudo-time analysis of keratinocytes subsets. a**, UMAP plot of the keratinocyte's subsets of scRNAseq aggregate performed on natural human skin modules after 0, 3, 7 or 10 days of culture, colored by cell type (left panel), and the associated pseudo-time visualization (right panel). **b**, Pseudo-time heatmap of gene expression along keratinocytes differentiation. **c**, Expression of undifferentiated (KRT14, ITGA6, ITGB1) and differentiated (KRT1, KRT10, DSG1) specific markers along the pseudo-time trajectory.

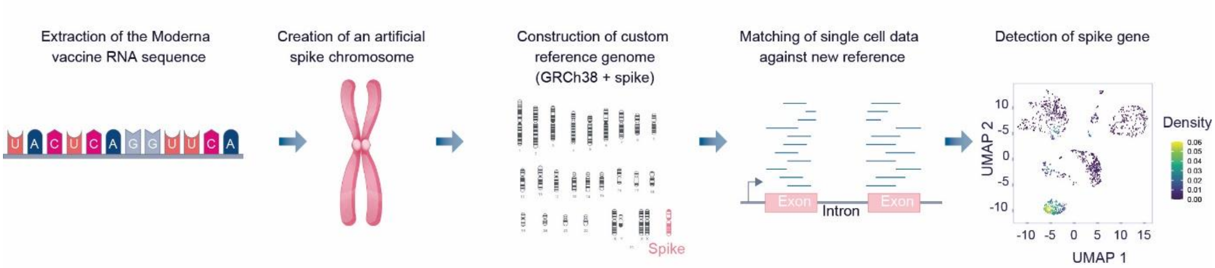


**Figure S4. Single cell RNA seq exogenous vaccine mRNA detection strategy.** First, the retro-engineered sequence of the mRNA used in COVID19 vaccine was extracted from literature. The corresponding DNA sequence was added to the GRCh38 genome as an artificial vaccine 'chromosome' in order to obtain a custom genome reference. The single cell raw data were matched against the custom genome reference, thus allowing the detection of a "spike gene" in the subsequently obtained count matrix.

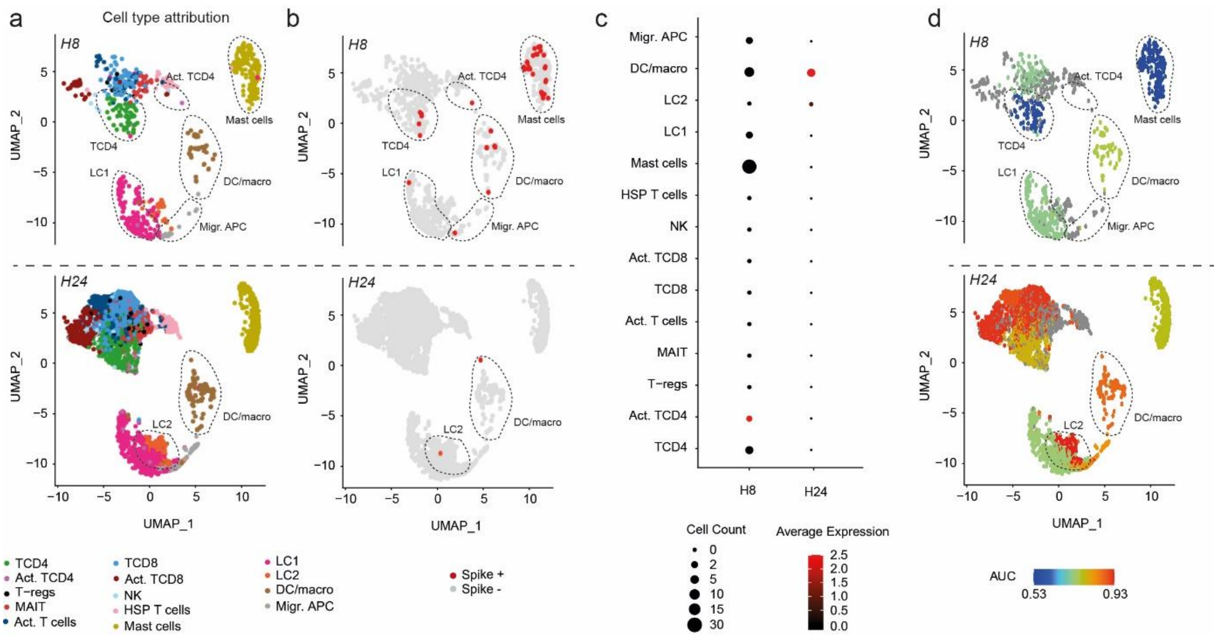


**Fig. S5. Identification of skin resident immune cells involved in internalization and response to i.d. injection of mRNA-1273 COVID-19 vaccine using scRNAseq. A**, UMAP of cell attribution at 8 (top) and 24 hours (bottom) after i.d. injection. **B**, UMAP plots indicating spike RNA detection in each cluster. **C**, Number of cells by cell-type which have incorporated spike mRNA and levels of expression. **d**, Analysis of gene expression perturbation due to COVID-19 vaccine i.d. injection with Augur. The AUC allows to evaluate the level of gene expression perturbation by cell type (AUC=0.5 no perturbation between conditions, the most the AUC is closed to 1, the most gene expression is different within a cell type between both conditions). *Act. TCD4: Activated CD4 T cells; Act. TCD8: Activated CD8 T cells; Act. T cells: Activated T cells; TCD4: CD4 T cells; TCD8: CD8 T cells; DC/macro: Dendritic Cells/macrophages; HSP T cells; T cells expressing Heat Shock Proteins; LC1: Langerhans Cells type 1; LC2: Langerhans Cells type 2; MAIT: Mucosal-Associated Invariant T cells; Migr. APC: Migratory Antigen Presenting Cells; NK: Natural Killer cells; T-regs: regulatory T cells.*

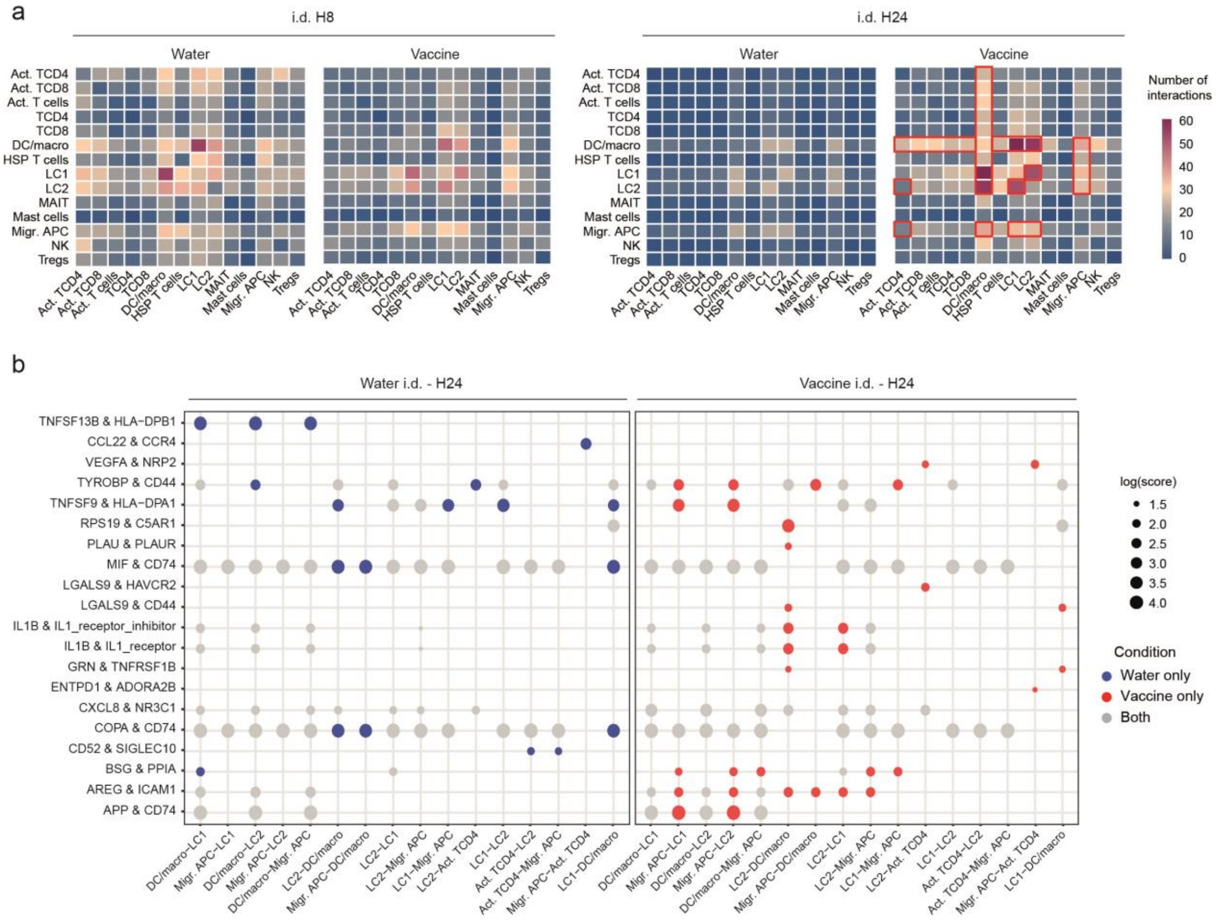


**Figure S6**. **Analysis of interactome between immune cell types after i.d. injection. a**, Analysis of the number of interactions between immune cell types 8 and 24 hours after i.d. injection of water or COVID-19 vaccine. **b**, Dot plot of the ligand-receptor interaction scores between immune cell types most affected by vaccine injection 24h after i.d. injection. Only statistically significant scores are presented.

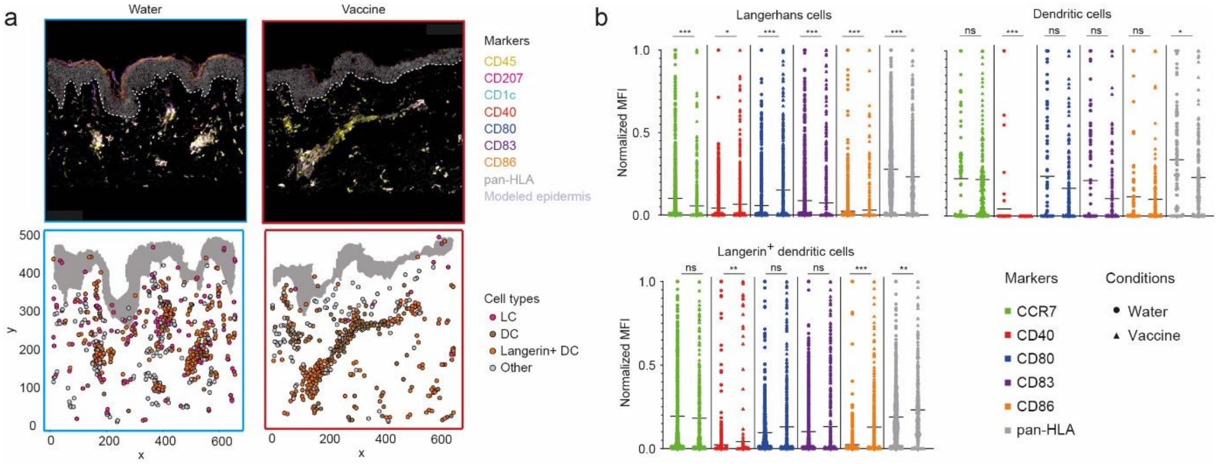


**Figure S7. Cellular activation of APC 8 hours post-vaccination. a**, Representative 3D confocal multiplex images (top) and associated digital maps (bottom) of MANTIS APC panel of subcutaneous water- (left) and vaccine-injected (right) human skin modules, 8 hours post-injection. **b**, Dotplot of activation markers (CCR7, CD40, CD80, CD83, CD86 and pan-HLA) normalized MFI in LCs (top left), DCs (top right) and Langerin+ DCs (bottom left) in water- (left) and vaccine-injected (right) human skin modules, 8 hours post-injection.
